# Differential ripple propagation along the hippocampal longitudinal axis

**DOI:** 10.1101/2022.12.22.521603

**Authors:** Roberto De Filippo, Dietmar Schmitz

## Abstract

Hippocampal ripples are highly synchronous neural events critical for memory consolidation and retrieval. A minority of strong ripples has been shown to be of particular importance in situations of increased memory demands. The propagation dynamics of strong ripples inside the hippocampal formation are, however, still opaque. We analyzed ripple propagation within the hippocampal formation in a large open access dataset comprising 267 Neuropixel recordings in 49 awake, head-fixed mice. Surprisingly, strong ripples (top 10% in ripple strength) propagate differentially depending on their generation point along the hippocampal longitudinal axis. The septal hippocampal pole is able to generate longer ripples that engage more neurons and elicit spiking activity for an extended time even at considerable distances. Accordingly, a substantial portion of the variance in strong ripple duration (R² = 0.463) is explained by the ripple generation location on the longitudinal axis. Our results are consistent with a possible distinctive role of the hippocampal septal pole in conditions of high memory demand.

## Introduction

Hippocampal ripples are brief oscillatory events detected in the local field potential (LFP) of the hippocampal formation, these events correspond to the synchronized depolarization of a substantial number of neurons in various hippocampal subregions (Hulse et al., 2016, Ylinen et al., 1995). An higher ripple incidence during memory encoding is associated with superior recall performance (Norman et al., 2019), furthermore, ripple incidence is increased during successful memory retrieval (Vaz et al., 2019, Carr et al., 2011). Ripples are also involved in memory consolidation both in awake and sleep conditions (Jadhav et al., 2012, Roux et al., 2017, Sirota et al., 2003, Girardeau et al., 2009), disrupting awake ripples during learning causes a persisting performance degradation, the same effect can be achieved by silencing ripples during post-learning sleep. Accordingly, ripples are considered to play a crucial role in memory processes and reorganization of memory engrams (Girardeau and Zugaro, 2011, Buzsáki, 2015, Diba and Buzsáki, 2007, Foster and Wilson, 2006, Xu et al., 2019, Takahashi, 2015, Davidson et al., 2009, Pfeiffer and Foster, 2015, Dragoi and Tonegawa, 2011, Girardeau et al., 2009). Ripples duration exhibits a skewed distribution with only a minority of long-duration ripples (> 100 ms). The fraction of long-duration ripples, ripple amplitude and within-ripple firing rate of both excitatory and inhibitory neurons are increased in both novel contexts and memory-demanding tasks (Fernández-Ruiz et al., 2019). Reducing ripple duration artificially causes a degraded working memory performance (Jadhav et al., 2012, Fernández-Ruiz et al., 2019) and, on the contrary, prolongation induced by optogenetic-activation has a beneficial effect (Fernández-Ruiz et al., 2019). Importantly, the artificial recruitment of additional neurons seems to be constrained by pre-existing resting potential dynamics (Noguchi et al., 2022). Hippocampal-neocortical interactions, suggested to be important for memory consolidation (Klinzing et al., 2019, Gais et al., 2007, Tukker et al., 2020), are increased specifically during long-duration compared to short-duration ripples (Ngo et al., 2020).

Ripple amplitude and duration are significantly correlated (Tong et al., 2021, Patel et al., 2013), moreover, they are both related to the amount of underlying spiking activity (Tong et al., 2021, Khodagholy et al., 2017). It is possible to combine ripple strength and amplitude by considering the area of the high-pass filtered envelope (‘ripple strength’).

These results point at a specific role of strong ripples (ripples with high strength) in situations of high mnemonic demand and are consistent with a possible power law distribution where a minority of ripples is responsible for a substantial part of memory requirements. For this reason, it is of interest to identify the possible electrophysiological peculiarities of this subgroup of ripples. Do strong ripples propagate differently compared to common ripples? Are strong ripples generated homogeneously along the hippocampal longitudinal axis? Do ripples have a preferred longitudinal directionality? In this study we focused our attention on ripples generation and propagation within the hippocampal formation. Hippocampal connectivity with cortical and sub-cortical areas varies considerably along the longitudinal axis (Moser and Moser, 1998, Fanselow and Dong, 2010) and gene expression, as well, exhibits both gradual and discrete transitions along the same axis (Vogel et al., 2020, Strange et al., 2014). Consequently, the hippocampus is considered to be functionally segmented along its long axis. The different connectivity contributes to explain the functional organization gradient between a predominantly spatio-visual (septal pole) and emotional (temporal pole) processing. Ripples generated in the septal and temporal hippocampal pole have already been shown to be temporally independent and able to engage different neuron subpopulations, even in the same downstream brain area (Sosa et al., 2020). Consequentially, a heterogeneous ripple generation chance along the longitudinal axis most probably has an impact on the frequency with which different brain areas and neurons subgroups are activated by ripples. Our work is based on a dataset provided by the Allen Institute (Siegle et al., 2021), this dataset enabled us to study comprehensively ripples features across the septal half of the hippocampus. Previous studies have looked at ripple propagation along the longitudinal axes of the hippocampus (Patel et al., 2013, Kumar and Deshmukh, 2020), however, the size of this dataset made it possible to unveil propagation details previously overlooked.

## Results

### Distance explains most of the ripple strength correlation variability

We studied ripple propagation along the hippocampal longitudinal axis in an open-access dataset provided by the Allen Institute. We analyzed the LFP signals across the visual cortex, hippocampal formation and brain stem (Supplementary Figure 1) simultaneous to ripples detected in the CA1 of 49 animals (average session duration = 9877.4 ± 43.1 seconds, average ripple incidence during non-running epochs = 2.49 ± 0.12 per 10s). Ripples (n ripples = 120462) were detected on the CA1 channel with the strongest ripple activity. Ripple strength (∫Ripple) was calculated as the integral of the filtered LFP envelope between the start and end points for every detected ripple. Ripple strength and duration are highly correlated in each session (mean r = 0.87 ± 0.005, Supplementary Figure 2). Notably ripple strength correlates significantly better with the hippocampal population spiking rate on a ripple-to-ripple basis compared to ripple duration alone (p = 4.31e-11, Supplementary Figure 3). Clear ripples were observed uniquely in the hippocampal formation (CA1, CA2, CA3, DG, SUB, ProS). Likewise, ripple-induced voltage deflections (RIVD, integral of the unfiltered LFP envelope) were also noticeably stronger in hippocampal areas (Supplementary Figure 4B-F). Ripple strength was noticeably irregular in single sessions both across time and space, even within the CA1 region (Supplementary Figure 4C). We focused on the variability in ripple strength across pairs of CA1 recording locations with clear ripple activity (n CA1 pairs = 303, n sessions = 46). Correlation of ripple strength across different CA1 regions was highly variable (Figure 1A-B-C) with a lower and upper quartiles of 0.66 and 0.87 (mean = 0.76, SEM = 0.01). Distance between recording location could explain the majority (57.6%) of this variability (Figure 1B) with the top and bottom quartiles of ripple strength correlation showing significantly different average distances (Figure 1C-D). Given the correlation variability we asked how reliably a ripple can travel along the hippocampal longitudinal axis. To answer this question, we looked at ripples lag in sessions that included both long-distance (> 2126.66 µm) and short-distance (< 857.29 µm) CA1 recording pairs (n sessions = 32, n CA1 pairs = 64, Figure 1E). Reference for the lag analysis was always the most medial recording location in each pair. Almost half of the ripples in long-distance pairs (49.3 ± 2.2%) were detected in both locations (inside a 120 ms window centered on ripple start at the reference location). Unsurprisingly short-distance pairs showed a more reliable propagation (69.59 ± 3.51%). Moreover, lag between long-distance pairs had a much broader distribution (Figure 1F) and a significantly bigger absolute lag (Figure 1G). Neither high nor short-distance pairs showed clear directionality (lag long-distance = −1.14 ± 0.64 ms, lag short-distance = −0.5 ± 0.41 ms). Looking at the relationship between lag and ripple strength in long-distance pairs, however, an asymmetric distribution was apparent (Figure 1F top), suggestive of a possible interaction between these two variables: stronger ripples appear to be predominantly associated with positive lags (i.e. ripples moving medial→lateral). To further investigate this relationship we divided ripples into two groups: strong (top 10% ripple strength per session at the reference location) and common (remaining ripples). The septal half of the hippocampus was divided in three sections with equal number of recordings: medial, central and lateral (Supplementary Figure 5). Strong ripples identified in the medial section, in opposition to common ripples, showed a markedly positive lag (lag = 17.83 ± 1.02 ms) indicative of a preferred medial→lateral travelling direction (Figure 1H top). Surprisingly, the same was not true for strong ripples identified in the lateral section (lag = 3.62 ± 1.05 ms, Figure 1I). Strong and common ripples lags were significantly different between medial and lateral locations both in common and strong ripples. A biased direction of propagation can be explained by an unequal chance of ripple generation across space. We can assume that selecting strong ripples we are biasing our focus towards ripples whose generation point (seed) is situated nearby our reference location, this would contribute to explain the unbalanced lag. This notion would, however, fail to explain the different directionality we observed between strong ripples in medial and lateral locations. This hints at a more complex situation.

**Figure 1.**
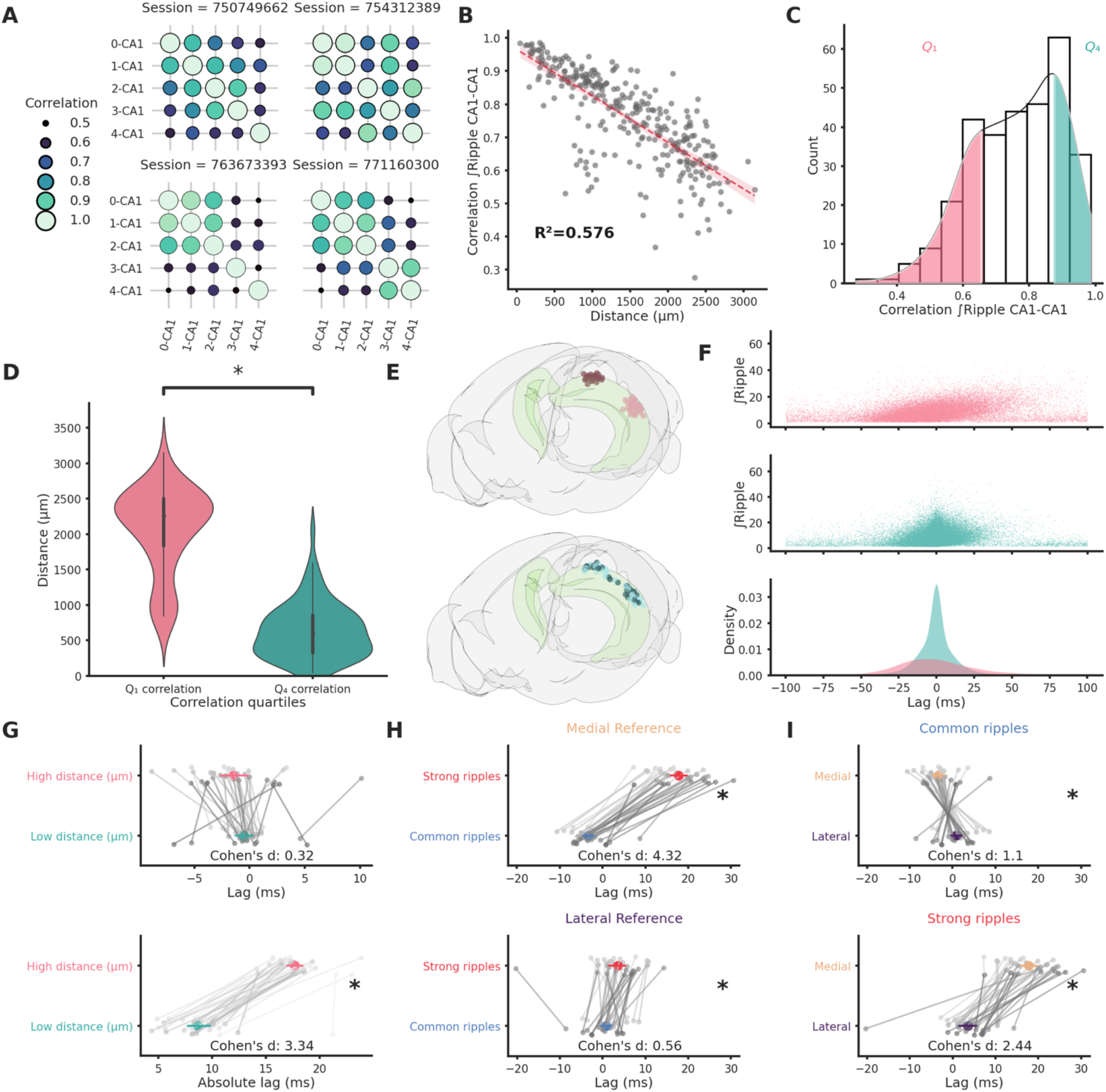
Ripple strength correlation depends significantly on distance. (A) Correlation matrices showing the variabilty of ripple strength correlation between pairs of recording sites located in different CA1 locations in 4 example sessions. The number on the x and y axis labels indicates the probe number. Probes are numbered according to the position on the hippocampal longitudinal axis (0 is the most medial probe). (B) Scatter plot and linear regression showing the relationship between distance and correlation strength. Distance between recording sites explains 0.576% of the variability in correlation of ripple strength. (C) Ripple strength correlation distribution. Pink represents bottom 25% (< Q₁) and blue top 25% (> Q₄). (D) Violinplots showing that the top and bottom correlation quartile show significantly different distance distributions (Q₁: 2077.57 ± 68.68 µm, Q₄: 633.56 ± 44.02 µm, p-value = 4.00e-23, Mann-Whitney U test). (E) Top: Rendering of the long distance (top) and short distance (bottom) CA1 pairs, dark circles are the reference locations in each pair. (F) Top and middle: scatter plots showing the relationship between ripple strength (at the reference location) and lag for long distance (top, n ripples = 31855) and short distance (middle, n ripples = 52858) pairs. Bottom: kernel density estimate of the lags of long distance (pink) and short distance (turquoise) pairs. (G): Lag (top) and absolute lag (bottom) comparison between long and short distance pairs (top: long distance =-1.47 ± 0.63 ms, Short distance = −0.51 ± 0.4 ms, p-value = 2.03e-01, Student’s t-test; bottom: long distance = 17.69 ± 0.38 ms, Short distance = 8.69 ± 0.56 ms, p-value = 6.58e-20, Student’s t-test). (H) Lag comparison in long distance pairs between common and strong ripples with reference located inthe medial (top) or lateral hippocampal section (bottom) (top: strong ripples=17.83 ± 1.02 ms, common ripples = −3.27 ± 0.68 ms, p-values = 2.28e-25, Student’s t-test, bottom: strong ripples=3.62 ± 1.05 ms, common ripples = 0.88 ± 0.66 ms, p-values = 3.00e-02, Student’s t-test). (I) Lag comparison in long distance pairs between ripples with reference located in the medial and lateral section in common (top) or strong ripples (bottom) (top: medial reference = −3.27 ± 0.68 ms, lateral reference = 0.88 ± 0.66 ms, p-values = 4.30e-05, Student’s t-test, bottom: strong ripples = 17.83 ± 1.02 ms, common ripples = 3.62 ± 1.05 ms, p-values = 4.30e-05, Student’s t-test).

### Ripples propagates differentially along the hippocampal longitudinal axis

To analyze the propagation of ripples along the hippocampal longitudinal axis we focused on sessions from which ripples were clearly detected in at least two different hippocampal sections at the same time (n = 41). We followed the propagation of strong and common ripples detected in the reference location across the hippocampus (Figure 2A-B) and built an average spatio-temporal propagation map per session (Figure 2C). Strong and common ripples in the medial section showed a divergent propagation pattern: strong ripples travelling medio→laterally and common ripples travelling in the opposite direction (Figure 2D-E). Ripples detected in the lateral section did not show such strikingly divergent propagation (Figure 2F-G) whereas, in the central section, the propagation was divergent only laterally and not medially (Figure 2H-I). This peculiar propagation profile suggests a not previously described underlying directionality along the hippocampal longitudinal axis and can be possibly explained by a spatial bias in strong ripples generation. To understand the mechanism underlying such difference in propagation we examined the location of the seed for each ripple in sessions in which ripples were clearly detected in every hippocampal section (n sessions = 25). While we found no differences in the number of ripples detected in each hippocampal section (p-value = 0.55, Kruskal-Wallis test), we observed differences regarding ripple generation. In common ripples, regardless of the reference location, most ripples started from the lateral section (Figure 3A left). On the other hand, strong ripples displayed a more heterogenous picture (Figure 3A right). We identified two principles relative to strong ripples generation: In all hippocampal sections the majority of strong ripples are locally generated, and a greater number of strong ripples is generated medially than laterally. Looking at the central section we can appreciate the difference between the number of strong ripples generated medially and laterally (Figure 3A right, mean medial = 36.83 ± 2.66%, mean lateral = 20.55 ± 2.04%, p-value = 3e-05, Pairwise Tukey test). Strong and common ripples had significantly different seed location profiles only in the medial and central section, not in the lateral section (Figure 3B). These seed location profiles contribute to explain the propagation idiosyncrasies: major unbalances in seeds location cause propagation patterns with clear directionality, on the contrary, lag measurements hovering around zero are the result of averaging between two similarly numbered groups of ripples with opposite direction of propagation. Notably, propagation speed did not change depending on the seed location (Supplementary Figure 6). The reason why strong ripples are only in a minority of cases generated in the lateral section remains nevertheless unclear. Using a ‘strength conservation index’ (SCI) we measured the ability of a ripple to retain its strength during propagation (a ripple with SCI = 1 is in the top 10% in all hippocampal sections). We observed that ripples generated laterally were effectively less able to retain their strength propagating towards the medial pole (Supplementary Figure 7). This result is not simply explained by differences in ripple strength along the medio-lateral (M-L) axis, as no such gradient was observed (R² = 0.0012, Supplementary Figure 8). Curiously, ripple amplitude showed a weak trend in the opposite direction (r = 0.25, p-value = 7.21e-04), with higher amplitude ripples in the lateral section (Supplementary Figure 9).

**Figure 2.**
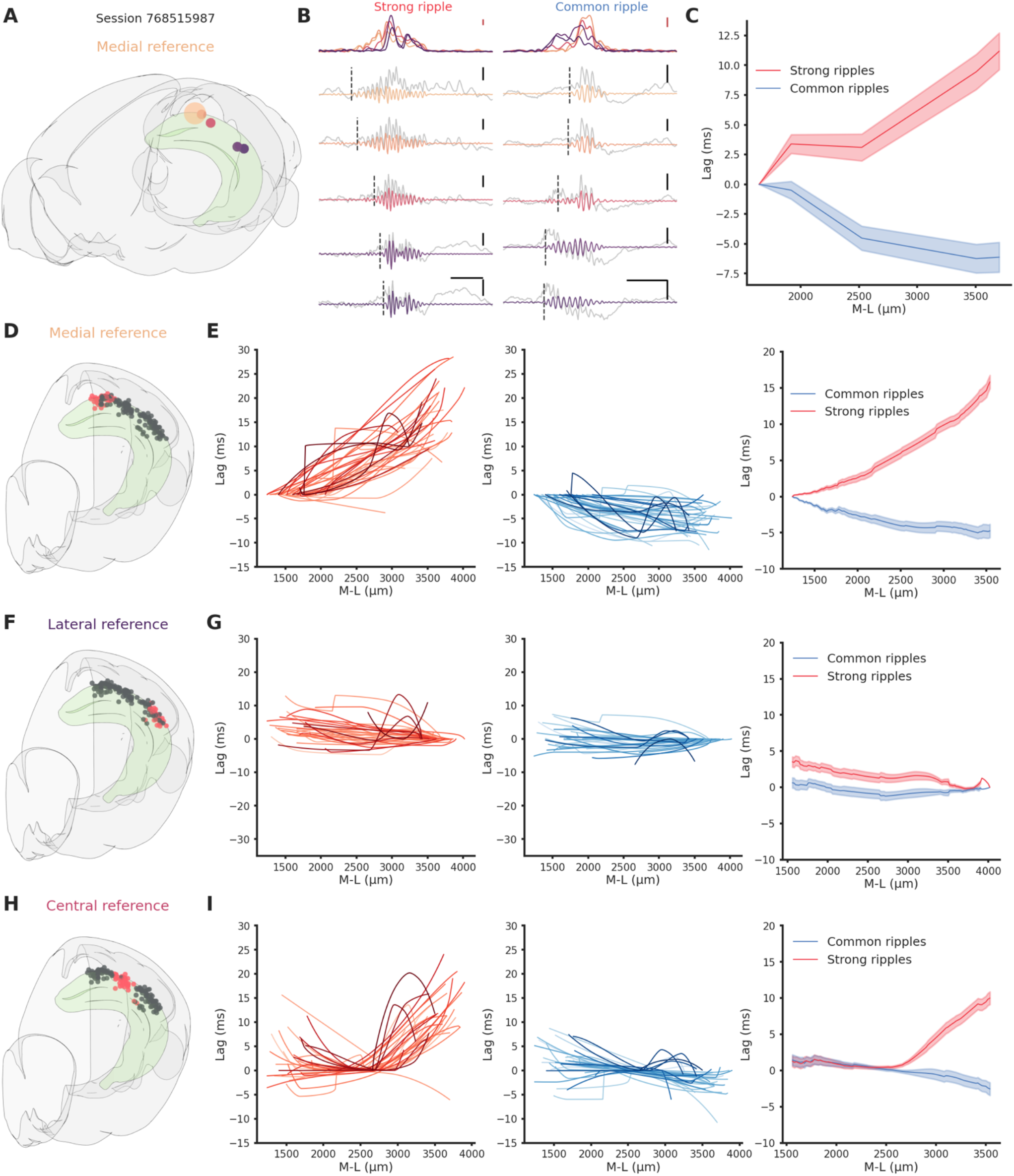
Direction-dependent differences in ripple propagation along the hippocampal longitudinal axis. (A) Recording locations for session 768515987. Circles colors represents medio-lateral location. Bigger circle represents the reference location. (B) Example propagation of a strong (left column) and common (right column) ripple across the different recording location from session 768515987, each filtered ripple is color-coded according to A. Grey traces represents raw LFP signal. Dashed vertical line represents the start of the ripple. In the top row the ripple envelope across all locations. Black scale bars: 50 ms, 0.5 mV. Red scale bars: 0.1 mV. (C) Average propagation map of strong and common ripples in session 768515987 across the medio-lateral axis. (D) Recording locations relative to E. Red circles represents the reference locations across all sessions (n sessions=41), black circles represents the remaining recording locations. (E) Left: Medio-lateral propagation of strong ripples, each line represents the average of one session. Middle: Medio-lateral propagation of common ripples, each line represents the average of one session. Right: Average propagation map across sessions of strong and common ripples. Reference locations are the most lateral per session. (F) Same as D. (G) Same as E. Reference locations are the most lateral per session. (H) Same as D. (I) Same as E. Reference locations are the most central per session.

**Figure 3.**
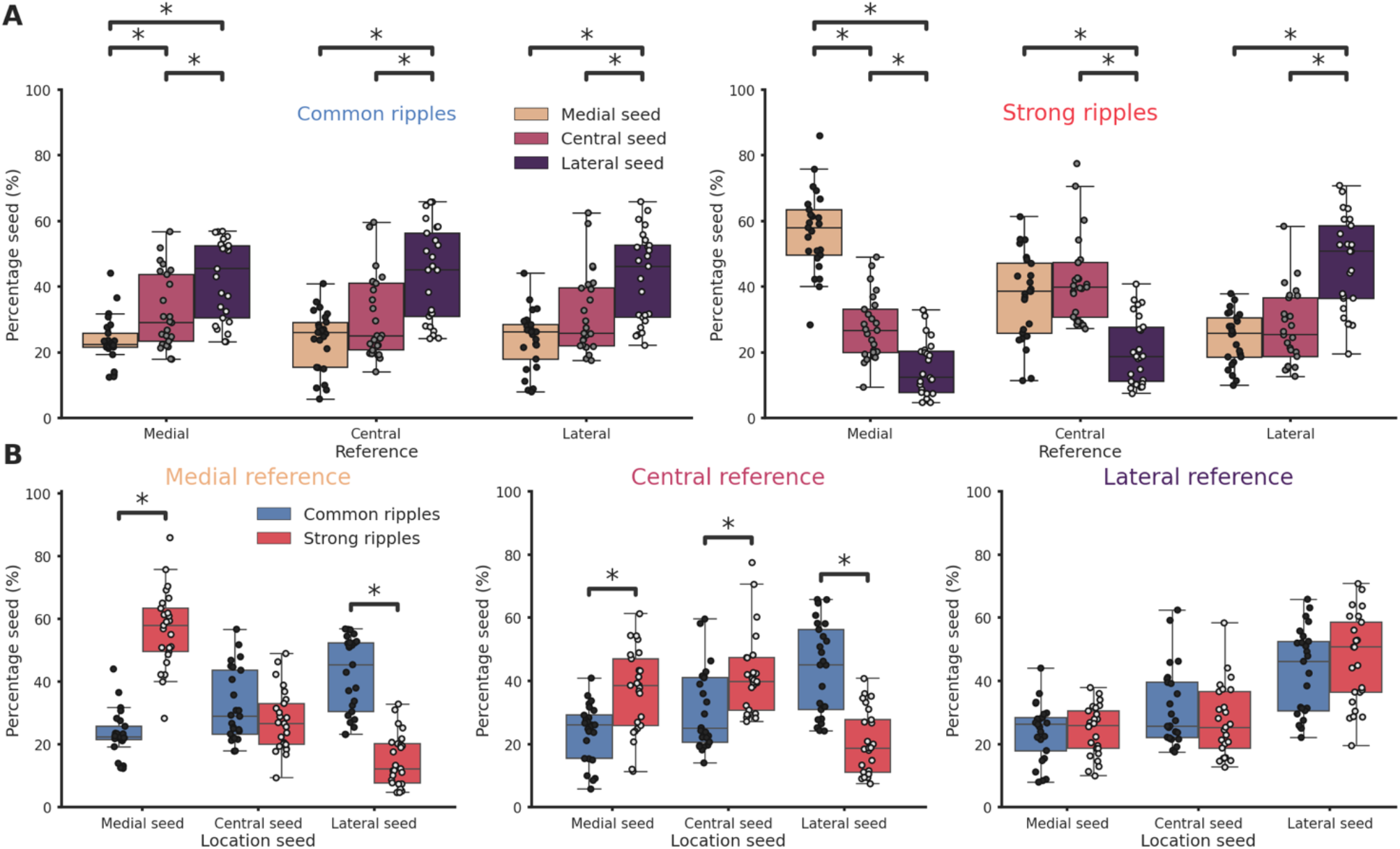
Ripples generation differences along the hippocampal longitudinal axis. (A) Ripple seed location comparison between the three reference locations in common ripples (left) and strong ripples (right). Majority of common ripples seeds are located in the lateral hippocampal section regardless of the reference location (medial reference/lateral seed = 42.43 ± 2.45 %, central reference/lateral seed = 43.77 ± 2.9 %, lateral reference/lateral seed = 42.83 ± 2.75 %). Strong ripples are mainly local (medial reference/medial seed = 56.78 ± 2.48 %, central reference/central seed = 41.74 ± 2.58 %, lateral reference/lateral seed = 46.76 ± 2.89 %).(B) Ripple seed location comparison between strong and common ripples using a medial (left), central (center) or lateral reference (right). Asterisks mean p < 0.05, Kruskal-Wallis test with pairwise Mann-Whitney post-hoc test.

### The hippocampal medial pole can generate longer ripples able to better engage neural networks

To understand the reason behind the differential propagation we focused uniquely on the central section, here it was possible to distinguish between ripples generated laterally or medially (‘lateral ripples’ and ‘medial ripples’). We included in the analysis sessions in which ripples were clearly detected in each hippocampal section and with at least 100 ripples of each kind (n sessions = 24). We looked at spiking activity associated with these two classes of ripples in the hippocampal formation across the M-L axis (n clusters per session = 650.42 ± 33.16, Figure 4A-B-C). To compare sessions, we created interpolated maps of the difference between spiking induced by medial and lateral ripples (Figure 4D). Immediately following ripple start (0-50 ms, “early phase”) spiking was predictably influenced by ripple seed proximity: in the lateral section lateral ripples induced more spiking (indicated by the blue color), whereas in the medial section medial ripples dominated (indicated by the red color). Surprisingly, in the 50-120 ms window post ripple start (“late phase”), medial ripples could elicit significantly higher spiking activity than lateral ripples along the entire M-L axis (Figure 4E). Dividing clusters in putative excitatory and inhibitory using the waveform duration we observed the same effect in both types of neurons (Supplementary Figure 10). In accordance with this result, we found that the medial hippocampal section is able to generate longer ripples (Figure 4F). An important portion of the variance in ripple duration is indeed explained by location on the M-L axis both in common (R² = 0.133) and especially in strong ripples (R² = 0.463). The observed extended spiking could be due to a increased number of neurons participating in the ripple, to a higher spiking rate per neuron or a combination of these two elements. Fraction of active neurons and spiking rate were both significantly higher in medial ripples (Supplementary Figure 11). Focusing only on the late phase the difference in fraction of active neurons per ripples between medial and lateral ripples was even more striking (Cohen’s d = 1.7, Figure 4G). Inversely, in the early phase, lateral ripples could engage more neurons, although, the effect size was much smaller (Cohen’s d = 0.39). The same result was found in relation to the spiking rate, medial ripples caused a significant and considerable increase in spiking rate in the late phase (Cohen’s d = 1.75, Figure 4H). Dividing again the clusters into putative excitatory and inhibitory, significant differences between medial and lateral ripples were present only in the late phase. Spiking frequency and number of engaged neurons were significantly higher in medial ripples both in putative excitatory and inhibitory clusters (Supplementary Figure 12). In summary, the prolonged spiking observed in medial ripples was caused both by an increased number of engaged neurons and a higher spiking rate per cell, both in putative excitatory and inhibitory neurons. The disparity in network engagement can possibly be in part explained by electrophysiological differences across hippocampal sections (e.g. higher firing rate). We did not find differences in the number of firing neurons (medial = 74.73, lateral = 79.8, p-value = 3.56e-01, Mann-Whitney U test), we did, however, found differences in firing rate, waveform duration, and waveform shape (recovery slope and peak-through ratio, Supplementary Figure 13). Firing rate and waveform duration exhibited respectively a left- and right-shifted distribution in the lateral section, reflecting lower firing rate and slower action potentials.

**Figure 4.**
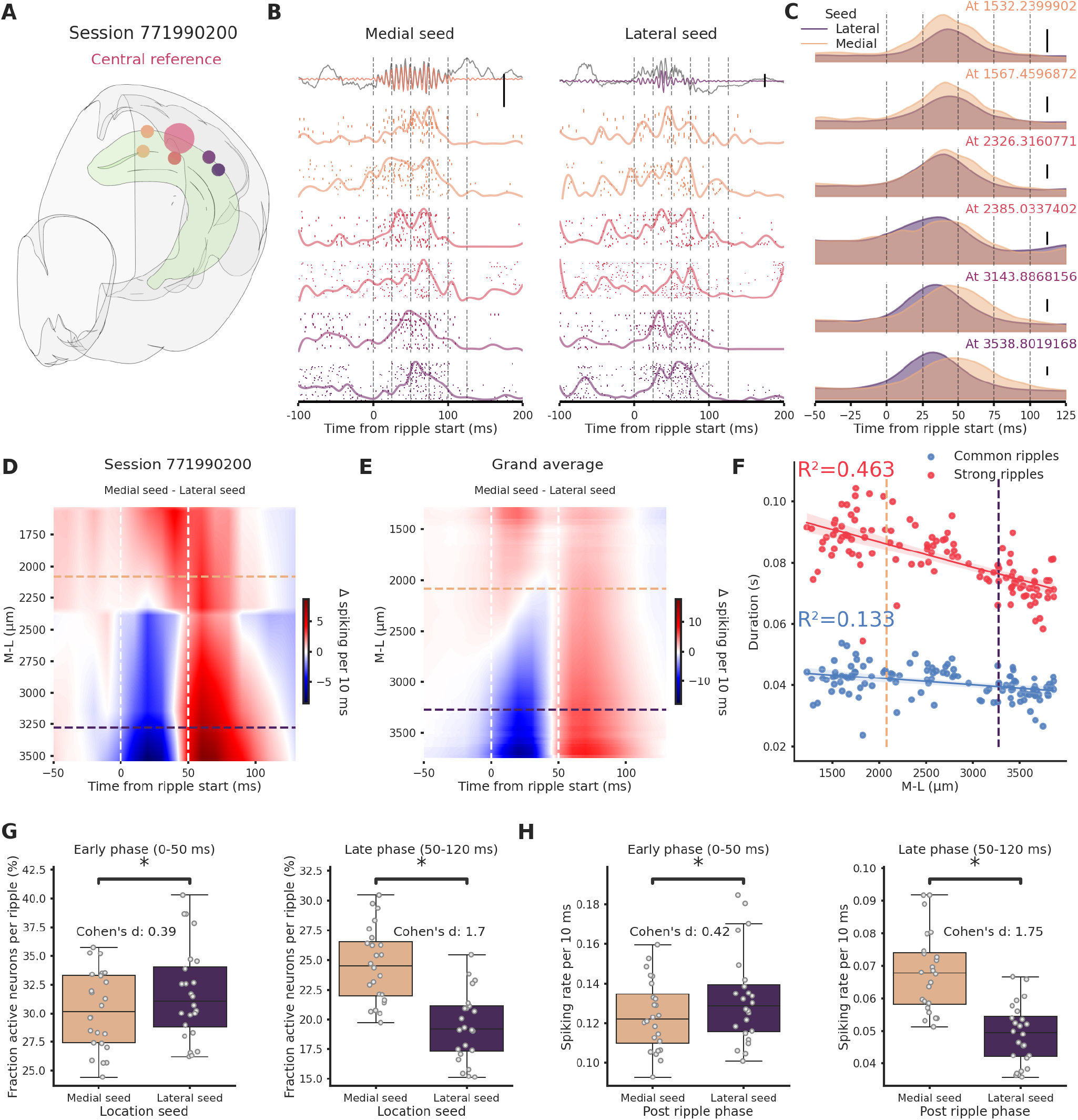
Ripples travelling in the medio→lateral direction show prolonged network engagement. (A) Recording location for session 771990200. Circles colors indicate medio-lateral location. Bigger circle represents the reference location. (B) Spiking activity across the hippocampal M-L axis associated with a ripple generated medially (left column) or lateraly (right column) across the different recording location from session 771990200. Spike raster plot and normalized density are plotted at each M-L location. In the top row filtered ripple, grey traces represents raw LFP signal. All plots are color coded according to A. Scale bar: 0.5 mV. (C) Kernel density estimates of the average spiking activity across different M-L locations and between seed type. Scale bar: 5 spikes per 10 ms. (D) Interpolated heatmap of the difference between medially and laterally generated ripple induced spiking activity in session 771990200. Vertical dashed lines represent borders between early and late post-ripple start phases. Horizontal dashed lines represent the spatial limits of the hippocampal sections. (E) Grand average of the differences between medially and laterally initiated ripple induced spiking activity across 24 sessions. Vertical dashed lines represent borders between early and late post-ripple start phases. Horizontal dashed lines represent the spatial limits of the hippocampal sections. (F) Regression plot between M-L location and ripple duration in common and strong ripples. Horizontal dashed lines represent the spatial limits of the hippocampal sections. (G) Average fraction of active neurons in medial (pink) and lateral (purple) ripples. Early/medial seed = 0.3 ± 0.69, early/lateral seed: 31.72 ± 0.84, p-value = 3.23e-05, Student’s t-test; late/medial seed = 24.57 ± 0.64, late/lateral seed = 19.44 ± 0.58, p-value = 4.09e-07, Student’s t-test. (H) Average spiking rate medial (pink) and lateral (purple) ripples. Early/medial seed = 0.12 ± 0.004, early/lateral seed = 0.13 ± 0.005, p-value = 1.35e-04, Student’s t-test; late/medial seed =0.07 ± 0.002, late/lateral seed = 0.05 ± 0.002, p-value = 1.24e-12, Student’s t-test.

## Discussion

Our results show for the first time that strong ripples propagate differentially along the hippocampal longitudinal axis. This propagation idiosyncrasy can be explained by a specific ability of the hippocampal septal pole (medial section in our analysis) to produce longer ripples that better entrain the hippocampal network and spread across the longitudinal axis. It was previously observed that ripples located at the septal and temporal pole are generated independently from each other, in addition, despite the presence of connections within the hippocampal longitudinal axis (Witter, 2007, van Strien et al., 2009), in the vast majority of cases ripples do not propagate to the opposite pole (Sosa et al., 2020). In accordance with these results, we observed a strong effect of spatial distance on ripple strength correlation confirming a previous study (Nitzan et al., 2022): the strength correlation, predictably, was higher in CA1 pairs closer to each other. The effect of distance was also apparent on the ripple chance of propagation, only half of the ripples generated in the septal pole were detected additionally in the intermediate hippocampus (lateral section in our analysis). This chance is much higher compared to the ∼3.7% reported regarding propagation between opposite poles (Sosa et al., 2020), it would be interesting to understand whether the temporal pole is also able to entrain the intermediate hippocampus in similar fashion or it is a peculiarity of the septal pole. A limitation of our work derives from the dataset being limited to the septal and intermediate hippocampus. Ripples can arise at any location along the hippocampal longitudinal axis (Patel et al., 2013). Our analysis shows that ripples are, however, not homogeneously generated across space. We observed important differences between strong ripples and common ripples generation. Common ripples followed a gradient with higher generation probability in the intermediate section and lowest in the septal pole. Strong ripples, on the other hand, were mostly generated locally (i.e. a strong ripple detected in the medial section is most likely generated in the medial section itself). Furthermore, only rarely a strong ripple generated in the intermediate hippocampus is able to propagate towards the septal pole retaining its strong status (top 10%). Conversely strong ripples generated in the septal pole have a significantly higher chance of propagate longitudinally and still be in the top 10% in terms of ripple strength. Notably, this is not consequence of a simple longitudinal gradient in ripple strength, indeed, we did not observe any difference in ripple strength along the longitudinal axis. Additionally, we show that ripples generated in the septal pole and in the intermediate hippocampus have a significantly different ability to engage hippocampal networks in the 50-120 ms window post ripple start. Ripples generated in the septal pole activate more neurons, both excitatory and inhibitory, and, moreover, elicit an higher spiking rate per neuron. This prolonged network activation is reflected by the fact that the position on the longitudinal axis explains 13.3% and 46.3% of the variability in ripple duration in common and strong ripples respectively. Consistent with a duration gradient along the longitudinal axis, the temporal hippocampus has been shown to produce shorter ripples both in awake and sleep conditions (Sosa et al., 2020). What is the reason that enables the septal pole to generate longer ripples? There might be for example underlying electrophysiological differences between the septal and intermediate hippocampus. Looking at units electrophysiological features we found some differences in the waveform shape and duration. We can hypothesize that slower action potentials and, consequentially, longer refractory periods hinder the ability to sustain protracted high frequency spiking. Accordingly, we found an increased firing rate and a smaller waveform duration in the septal pole. This might contribute to explain the prolonged ripples observed in the septal pole. We can also speculate that the neuromodulatory inputs gradient, monoamine fibers have been shown to be stronger in the ventral part (Strange et al., 2014), might influence neurons responses. Serotonin (ul Haq et al., 2016, Wang et al., 2015), noradrenaline (Ul Haq et al., 2012, Novitskaya et al., 2016) and acetylcholine (Zhang et al., 2021) have all been shown to suppress ripples. In accordance with this, some ripples are coupled with a reduced activation of the locus coeruleus and the dorsal raphe nucleus in vivo (Ramirez-Villegas et al., 2015).

Ripples can be subdivided in different types according to the relationship between the hippocampal LFP and the ripple itself (Ramirez-Villegas et al., 2015). Intriguingly these subtypes are associated with two different brain-wide networks, the first communicating preferentially with the associative neocortex and a second one biased towards subcortical structures. Moreover, these different types of ripples have been proposed to possibly fulfill different functional roles. Given the different input/output connectivity between septal, intermediate and temporal hippocampus (Fanselow and Dong, 2010) we hypothesize that ripple generated at different points of the hippocampal longitudinal axis might as well have functional differences, with the longer ripples generated septally possibly able to combine the different kind of informations processed in the distinct hippocampal sections and additionally relaying the integrated information back to the neocortex in accordance with the two-stage memory hypothesis (Diekelmann and Born, 2010, Marr, 1971, Buzsáki, 1989, Rasch and Born, 2007, McClelland et al., 1995). Long duration ripples have been shown to be of particular importance in situations of high-memory demand (Fernández-Ruiz et al., 2019), at the same time, previous studies highlighted the role of septal hippocampus in memory tasks and information processing (Hock and Bunsey, 1998, Moser et al., 1993, Moser et al., 1995, Steffenach et al., 2005, Kheirbek et al., 2013, McGlinchey and Aston-Jones, 2018, Fanselow and Dong, 2010, Maras et al., 2014, Bradfield et al., 2020, Qin et al., 2020). Our results can contribute to explain the specific role of septal hippocampus in memory-demanding tasks with its ability of generating particularly long ripples that are able to strongly engage networks in the entire top half of the hippocampal formation for an extended time.

## Materials and Methods

### Dataset

Our analysis was based on the Visual Coding - Neuropixels dataset provided by the Allen Institute and available at https://allensdk.readthedocs.io/en/latest/visual_coding_neuropixels.html. We excluded 6 sessions because of absence of recording electrodes in CA1 (session ids=732592105, 737581020, 739448407, 742951821, 760693773, 762120172). Furthermore, one session was excluded (session id = 743475441) because of an artifact in the LFP time series (time was not monotonically increasing) and two other sessions (session ids = 746083955, 756029989)because of duplicated LFP traces (see https://github.com/RobertoDF/Allen_visual_dataset_artifacts/blob/main/check_lfp_errors_from_files.ipynb). Our analysis was therefore focused on 49 sessions, average animal age = 119.22 ± 1.81. Sex: males n = 38, females n = 11. Genotypes: wt/wt n = 26, Sst-IRES-Cre/wt;Ai32(RCL-ChR2(H134R)_EYFP)/wt n = 10, Vip-IRES-Cre/wt;Ai32(RCL-ChR2(H134R)_EYFP)/wt n = 7, Pvalb-IRES-Cre/wt;Ai32(RCL-ChR2(H134R)_EYFP)/wt n = 6. Average probe count per session = 5.73 ± 0.08. Average number of recording channels per session = 2129.45 ± 29.46. Probes in each session were numbered according to the position on the M-L axis, with probe number 0 being the most medial. Channels with ambiguous area annotations were discarded (e.g. HPF instead of CA1). We found a number of of small artifacts in a variety of sessions, all this timepoints were excluded from the analysis (for more informations: https://github.com/RobertoDF/Allen_visual_dataset_artifacts). Further details about data acquisition can be found at https://brainmapportal-live-4cc80a57cd6e400d854-f7fdcae.divio-media.net/filer_public/80/75/8075a100-ca64-429a-b39a-569121b612b2/neuropixels_visual_coding_-_white_paper_v10.pdf. Visualization of recording locations was performed with brainrender (Claudi et al., 2021).

### Ripples detection

The LFP traces sampled at 1250 Hz were filtered using a 6th order Butterworth bandpass filter between 120.0 and 250.0. Ripples were detected on CA1 LFP traces, the best channel (higher ripple strength) was selected by looking at the SD of the envelope of the filtered trace, if multiple SD peaks were present across space (possibly caused by sharp waves in stratum radiatum and ripple activity in stratum pyramidale) we subsequently looked at the channel with higher skewness, in this way we could reliably identify the best ripple channel. The envelope of the filtered trace was calculated using the Hilbert transform (scipy.signal.hilbert). Ripple threshold was set at 5 SDs. Start and stop times were calculated using a 2 SDs threshold on the smoothed envelope with window = 5 (pandas.DataFrame.rolling) to account for ripple phase distortions. Ripple amplitude was calculated as the 90th percentile of the envelope.Ripple duration was limited at > 0.015 s and < 0.25 s. Candidate ripples with starting times closer than 0.05 s were joined in a single ripple with peak amplitude being the highest between the candidates. We estimated power density of each candidate using a periodogram with constant detrending (scipy.signal.periodogram) on the raw LFP trace, we checked the presence of a peak > 100 Hz, candidates not fulfilling this condition were discarded, this condition was meant to reduce the number of detected false positives. Ripple candidates detected during running epochs were discarded, an animal was considered to be running if his standardized speed was higher than the 10th percentile plus 0.06. Candidates were also discarded if no behavioral data was available. Code for the detection of ripples resides in ‘Calculate_ripples.py’.

### Correlation and lag analysis

In each session we uniquely used ripples from the CA1 channel with the strongest ripple activity, we looked at the LFP activity in all brain areas recorded in a window of 100.0 ms pre ripple start and 200.0 ms post ripple start, this broad windows account for possible travelling delays due to distance. For each brain area we picked the channel with higher SD of the envelope of the filtered trace. For each ripple considered we calculated integral of the envelope of the filtered trace (∫Ripple) and the integral of the raw LFP (ripple-induced voltage deflection, RIVD). After discarding channels with weak ripple activity (envelope variance < 5), we computed the pairwise pearson correlation of the envelope traces of CA1 channels (pandas.DataFrame.corr). For the lag analysis we first identified pairs of CA1 that satisfied a distance requirements. Distance threshold were set at 25% (857.29 µm) and 75% (2126.66 µm) of the totality of distances. For each ripple detected in the reference channel we identifired the nearest neighbour in the other channel. The analysis was repeated after dividing ripples in strong (top 10% ∫Ripple) and common ripples (all remaining ripples) per session. Code for the correlation and lag analysis resides in ‘Calculations_Figure_1.py’.

### Ripple spatio-temporal propagation maps and ripple seed analysis

The hippocampus was divided in three section with equal number of recordings. Channels with weak ripple activity (envelope variance < 5) were discarded. Sessions with recording locations only in one hippocampal sections or with less than 1000 ripples in the channel with strongest ripple activity were discarded as well. For each ripple detected on the reference CA1 channel we identified ripples in other CA1 channels happening in a window of ± 60.0 ms, this events were grouped together in a ‘cluster’. If more than one event was detected on the same probe we kept only the first event. ‘Clusters’ were subsequently divided according to ∫Ripple on the reference electrode in strong and common ripples. Lag maps were result of averaging lags for each probe. Code for the calculations of propagation maps resides in ‘Calculate_trajectories.py’.

### Ripple-associated spiking activity

We focused on sessions with clear ripple activity (envelope variance > 5) in all three hippocampal sections and at least 100 ripples generated both medially and laterally. The reference was always placed in the central section, here it was possible to identify ripples generated medially and laterally. We only considered ripples that were detected in at least half of the recording electrodes (in the code: “spatial engagment” > 0.5). For each ripple we computed a histogram of spiking activity of regions belonging to the hippocampal formation (HPF) in a window of 0.5 s centered on the ripple start in each probe. We averaged all the computed histograms to create a spatial profile of spiking activity. To compare spiking activity between sessions we interpolated (xarray.DataArray.interp) the difference between medial ripples-induced spiking and lateral ripples-induced spiking over space (this was necessary because probes in each sessions have different M-L coordinates) and time. We calculated the number of active cells (at least one spike) and spiking rate of each cluster per ripple in a window of 0.12 s starting from ripple start. We repeated the analysis separating the 0-50 ms and 50-120 ms post ripple start windows.

### Units selection and features calculations

Clusters were filtered according to the following parameters: Waveform peak-trough ratio < 5, ISI violations < 0.5, amplitude cutoff < 0.1 and Presence ratio > 0.1. For an explanation of the parameters see https://github.com/AllenInstitute/ecephys_spike_sorting/blob/master/ecephys_spike_sorting/modules/quality_metrics/README.md and https://brainmapportal-live-4cc80a57cd6e400d854-f7fdcae.divio-media.net/filer_public/80/75/8075a100-ca64-429a-b39a-569121b612b2/neuropixels_visual_coding_-_white_paper_v10.pdf. Firing rate was calculated on all clusters with presence ratio > 0.1.

## Acknowledgements

This study was supported by the German Research Foundation Deutsche Forschungsgemeinschaft (DFG), project 184695641 - SFB 958, project 327654276 - SFB 1315, Germany’s Excellence Strategy - Exc-2049-390688087 and by the European Research Council (ERC) under the European Union’s Horizon 2020 research and innovation programme (Grant agreement No. 810580). We thank J.T. Tukker, N. Maier for feedback on an early version of the manuscript and the members of the Schmitz lab for scientific discussion. We thank Willy Schiegel and Tiziano Zito for technical help with cluster computing. We thank Federico Claudi for support with brainrender. The authors declare that they have no competing interests.

## Contributions

Conceptualization, data curation, formal analysis, investigation, visualization: RDF. Writing - original draft: RDF. Writing - review & editing: RDF, DS. Funding acquisition: DS.

## Data and materials availability

All the code used to process the dataset is available at https://github.com/RobertoDF/De-Filippo-et-al-2022, pre-computed data structures can be downloaded at 10.6084/m9.figshare.20209913. All figures and text can be reproduced using code present in this repository, each number present in the text is directly linked to a python data structure. The original dataset is provided by the Allen Institute and available at https://allensdk.readthedocs.io/en/latest/visual_coding_neuropixels.html.

## Supplementary Figures

**Supplementary Figure 1.**
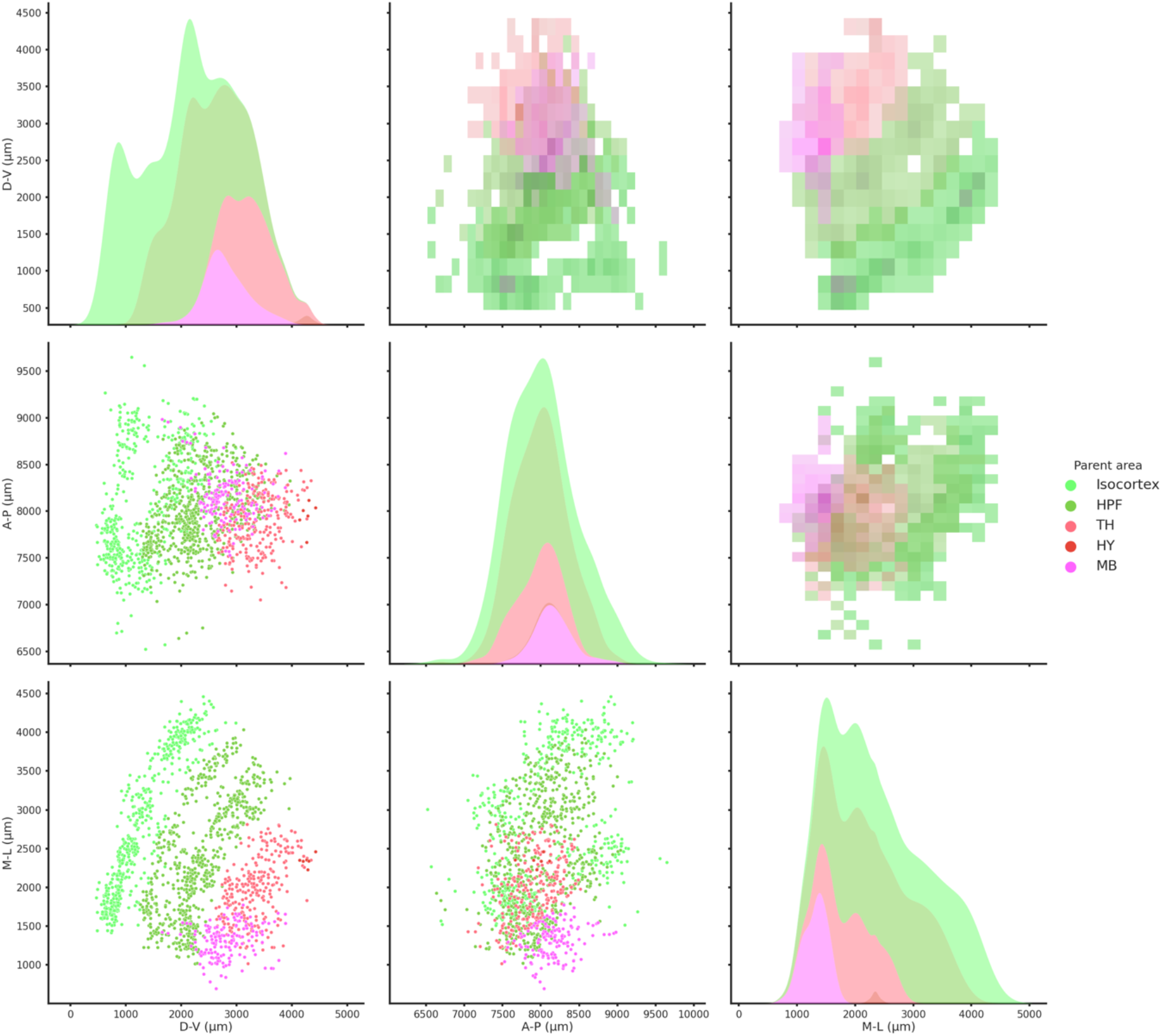
Spatial coordinates of all recorded brain regions. 2D histograms (upper diagonal), scatter plots (lower diagonal) and kernel density estimate plots (diagonal) of all the recorded regions color-coded according to the Allen Institute color scheme. HPF=hippocampus, TH=thalamus, HY=hypothalamus and MB=midbrain. M-L axis is zeroed at the midline.

**Supplementary Figure 2.**
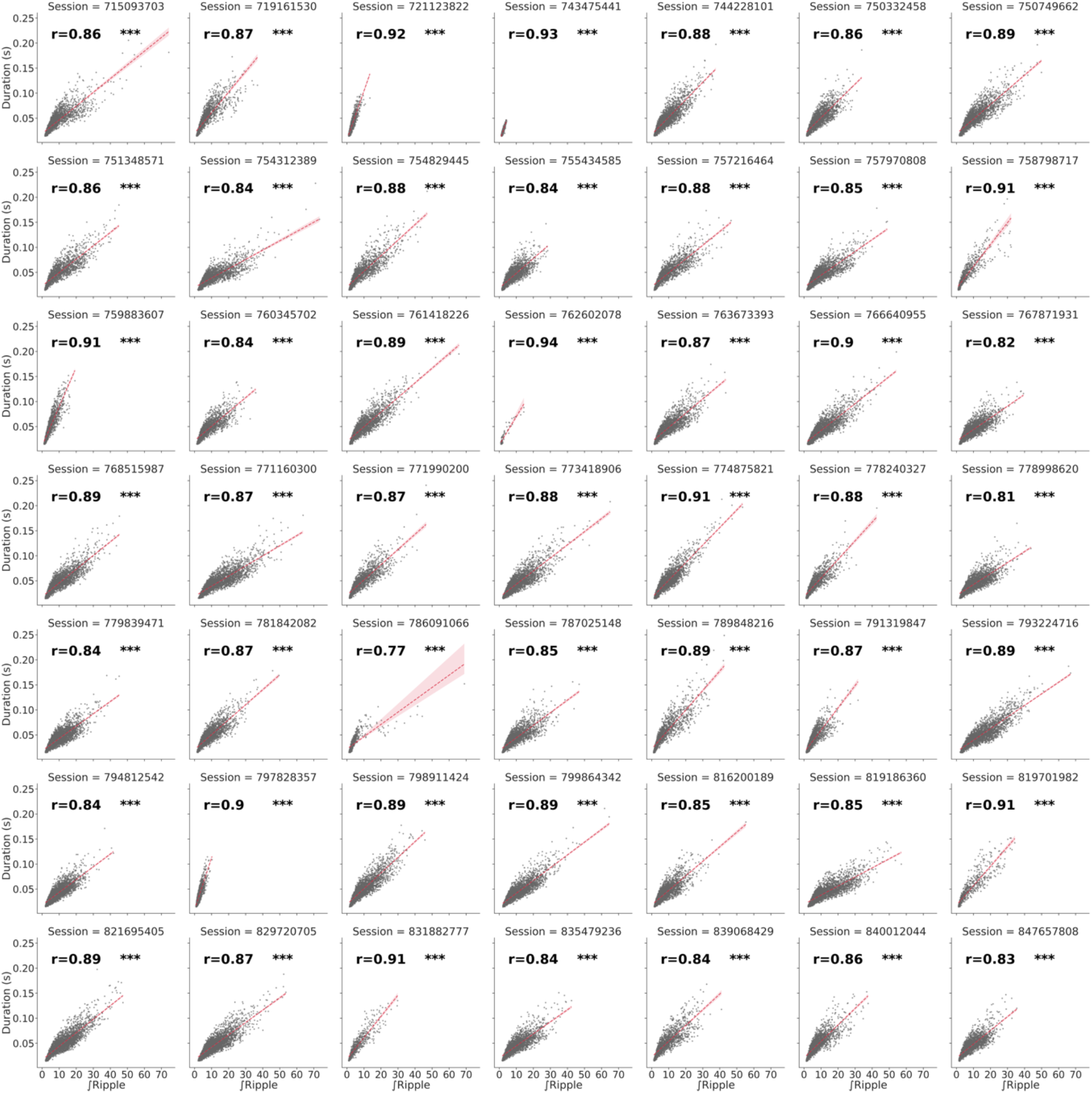
Correlation between ripple duration and strength per session. Red line represents linear regression with confidence interval of 95% estimated via bootstrap. *** means p < 0.0005.

**Supplementary Figure 3.**
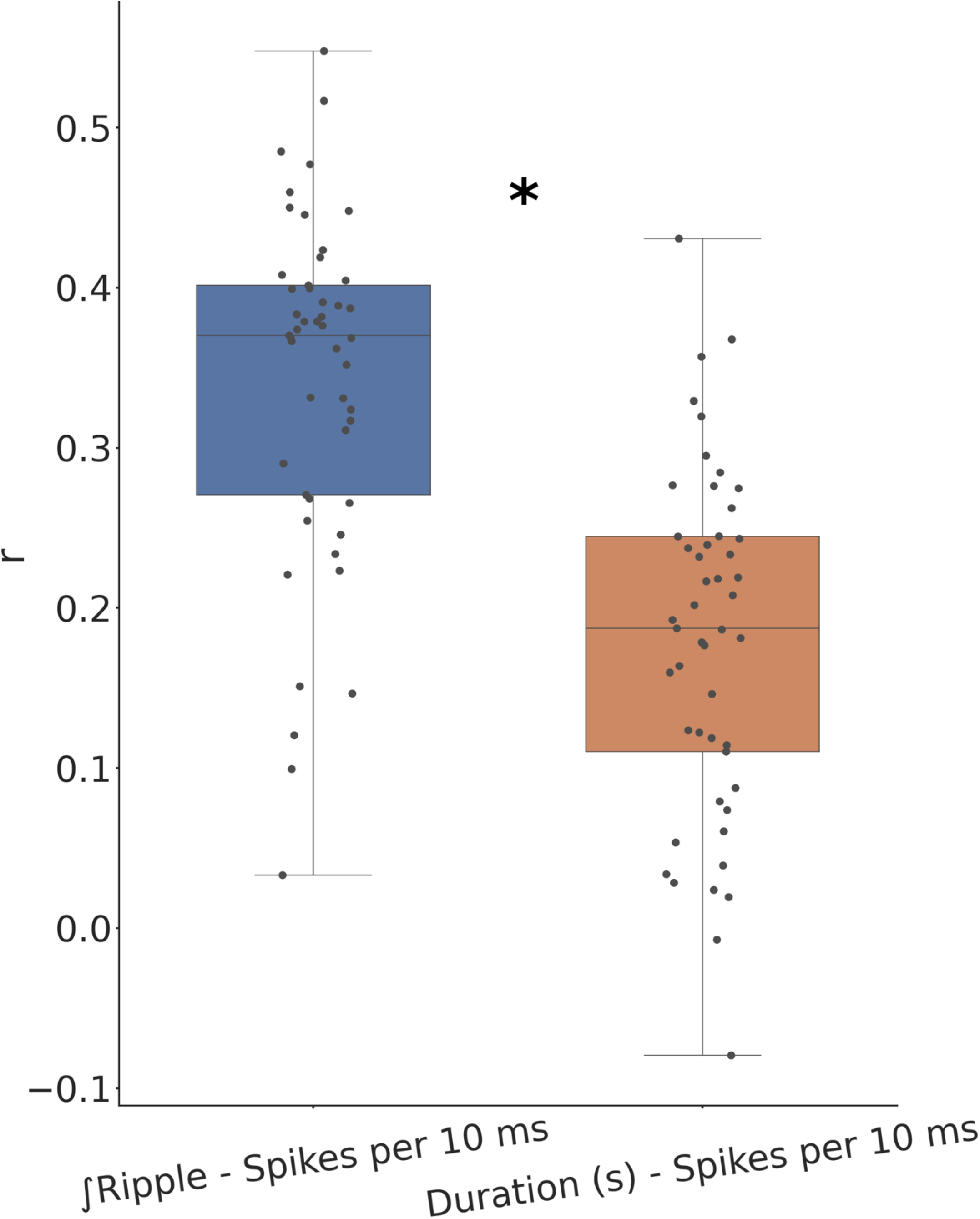
Comparison between correlation of ripple strength and duration with underlying spiking. Ripple strength correlates significantly better with the underlying ripple spiking activity. * means p < 0.0005.

**Supplementary Figure 4.**
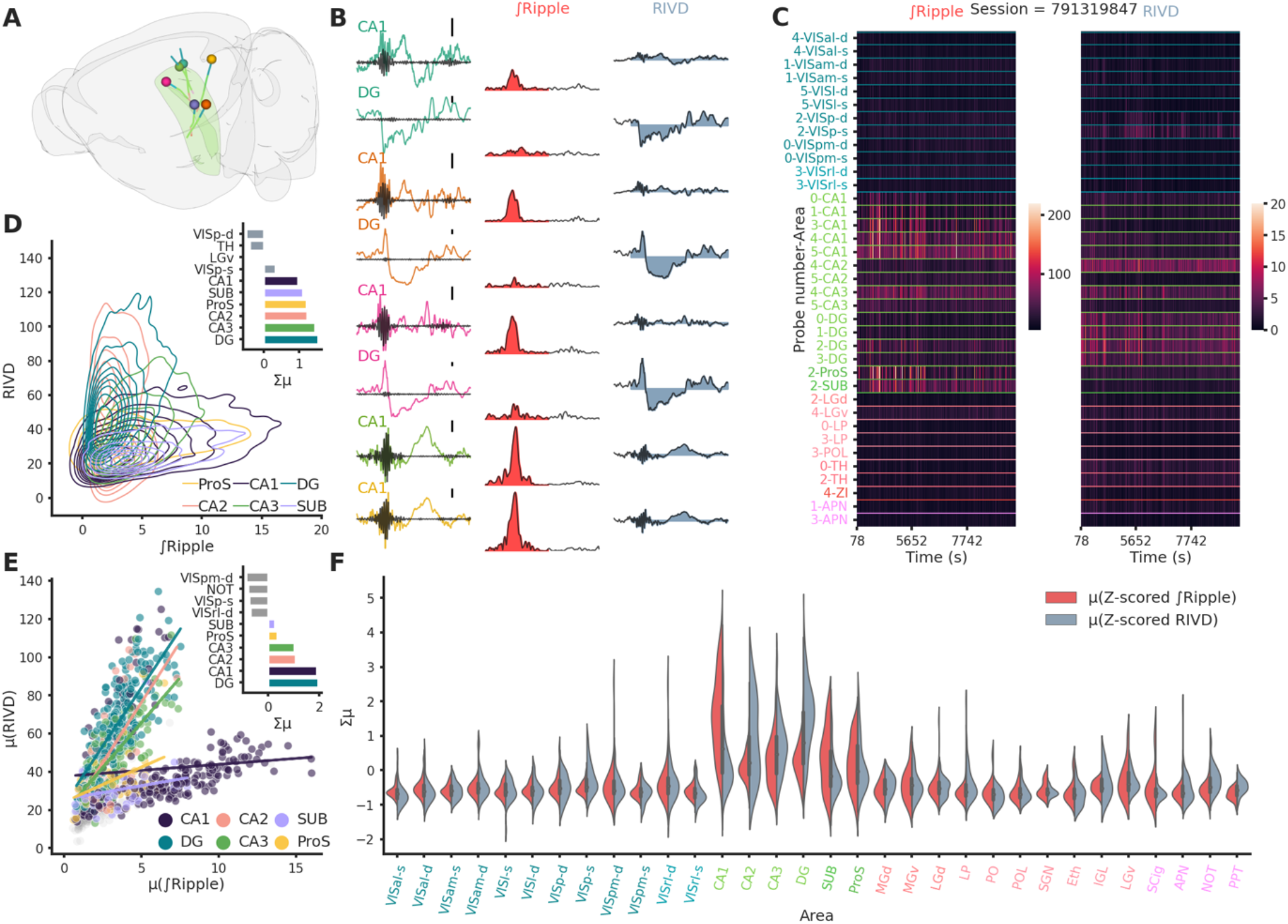
Ripple-associated LFP responses are predominantly observed in hippocampal structures. (A) Rendering of probe locatiosn for session 791319847. (B) First column: Raw LFP traces color coded according to probe identity, superimposed in black the trace after high-pass filtering to show the presence of a ripple. Scale bar: 250 µV. Middle column: Ripple envelope and associated ∫Ripple in red. Last column: Raw LFP trace and associated RIVD in blue. (C) Heatmaps of ∫Ripple (left) and RIVD (right) for the entirety of session 791319847 and for each recorded area. To note the variability in ∫Ripple over time and cross different CA1 locations.(D) Kernel density estimate plot showing the relationship between ∫Ripple and RIVD. Bar plot shows the sum of the z-scored ∫Ripple and RIVD per area.for the areas showing the strongest responses in session 791319847. (E) Summary scatter plot showing the relationship between ∫Ripple and RIVD for all sessions. Bar plot shows the sum of the z-scored ∫Ripple and RIVD per area averaged across animals. Most of the activity is confined to the hippocampal formation (DG, CA1, CA2, CA3 Sub and ProS) (n=49). (F) Violin plots showing the distribution of ∫Ripple and RIVD z-scored per session, hippocampal regions (text in green) show the biggest responses.

**Supplementary Figure 5.**
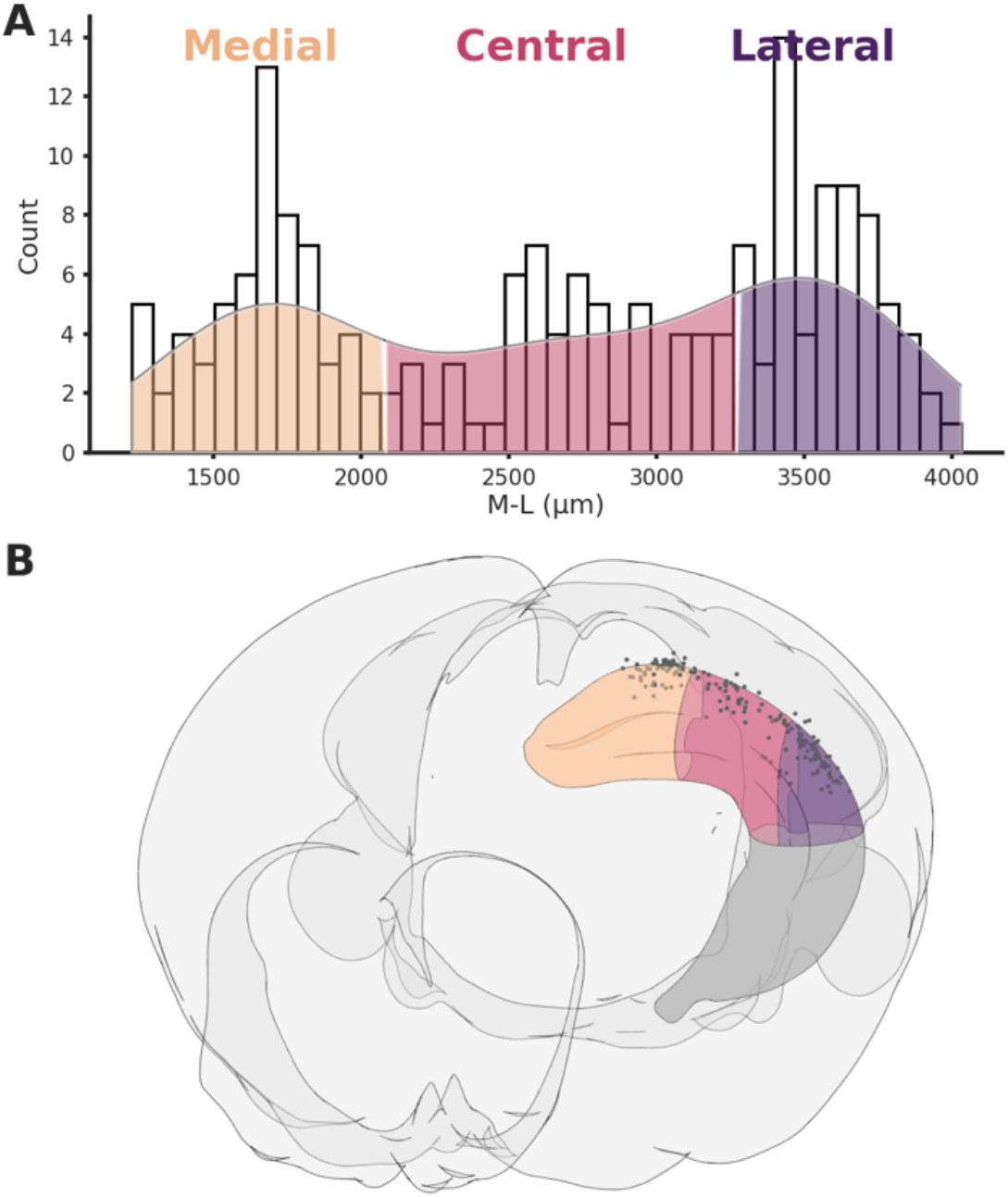
Hippocampal sections. (A) Histogram showing the three sections across the M-L axis, the hippocampus was divided in order to have an equal number of recordings in each section. (B) Rendering of the 3 sections and associated recording locations (black dots).

**Supplementary Figure 6.**
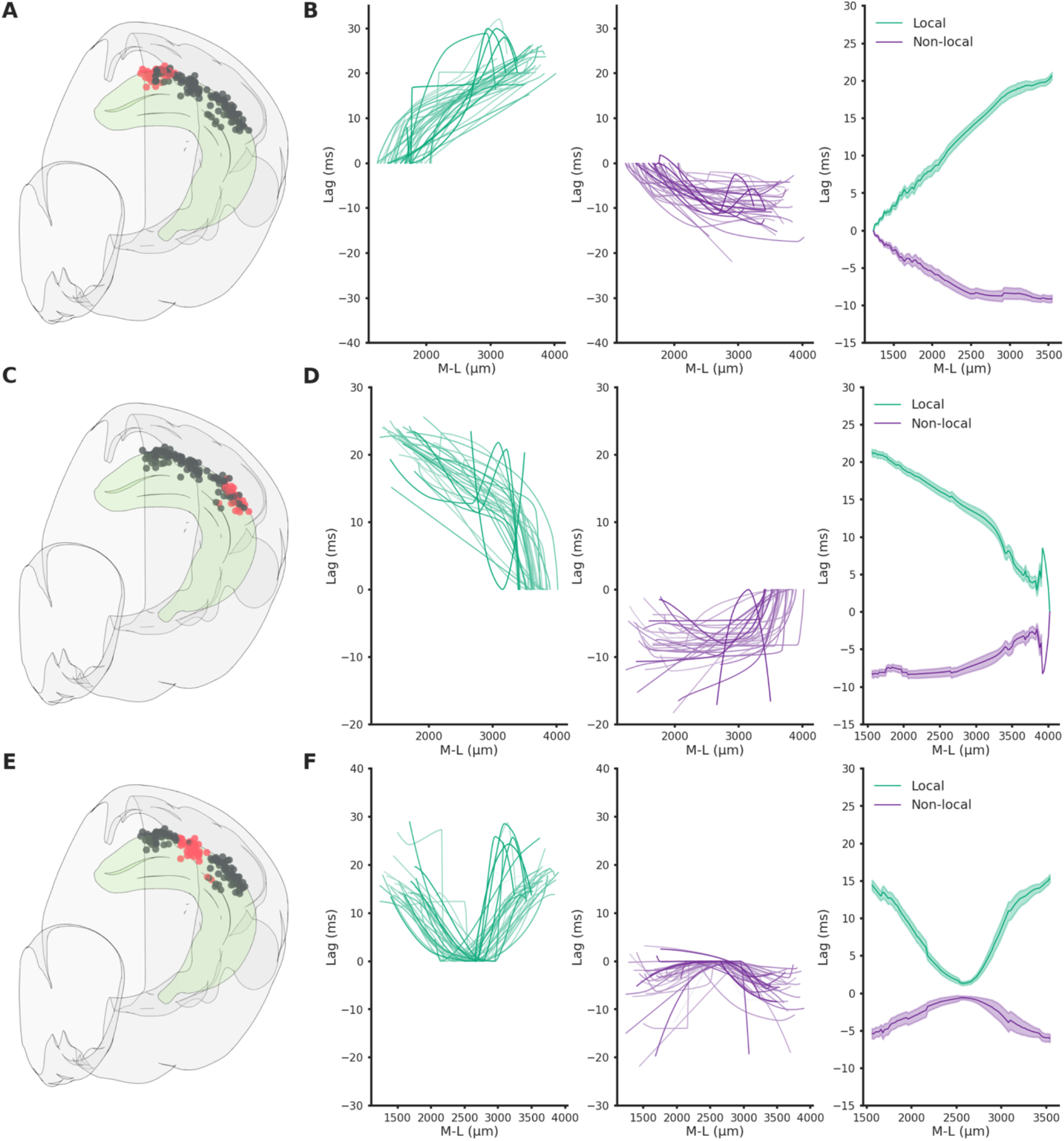
Spatio-temporal lag maps of locally and not locally generated ripples. Spatio-temporal profiles are symmetrical, strong indication of similar propagation speed regardless of seed position. (A) Recording locations relative to (B). Red circles represents the reference locations across all sessions (n sessions=41), black circles represents the remaining recording locations. (B) Left: Medio-lateral propagation of locally generated ripples (generated in the reference section), each line represents the average of one session. Middle: Medio-lateral propagation of non-locally generated ripples, each line represents the average of one session. Right: Average propagation map across sessions of strong and common ripples. Reference locations are the most lateral per session. (C) Same as A. (D) Same as B. Reference locations are the most lateral per session. (E) Same as A. (F) Same as B. Reference locations are the most central per session.

**Supplementary Figure 7.**
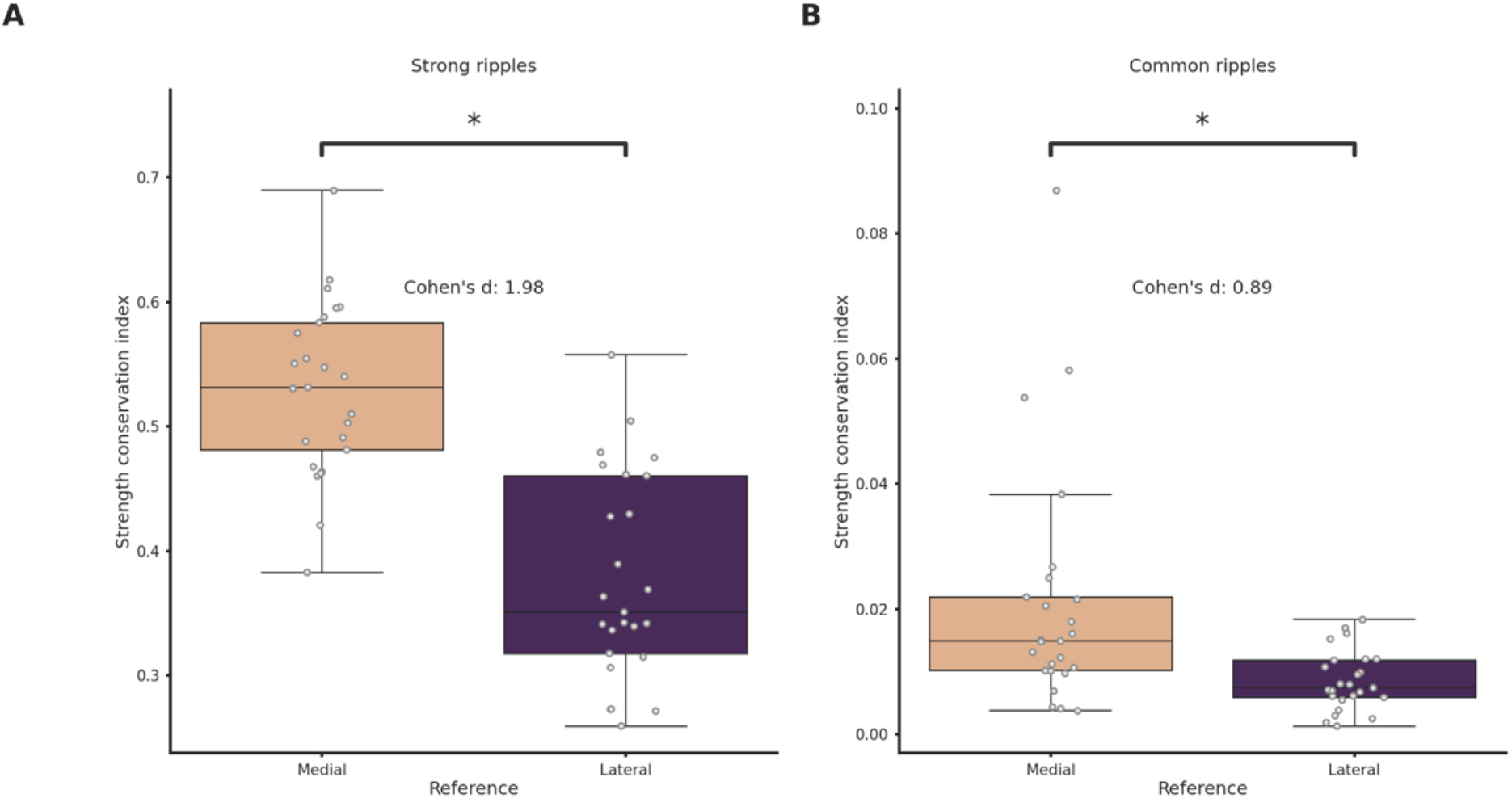
Strength conservation in medially and laterally generated ripples. (A) Strength conservation index in strong ripples grouped by reference location. Ripples generated in the lateral section showsignificantly lower strength conservation (p=7e-09, Student’s t-test). (B) Strength conservation index in common ripples grouped by reference location.

**Supplementary Figure 8.**
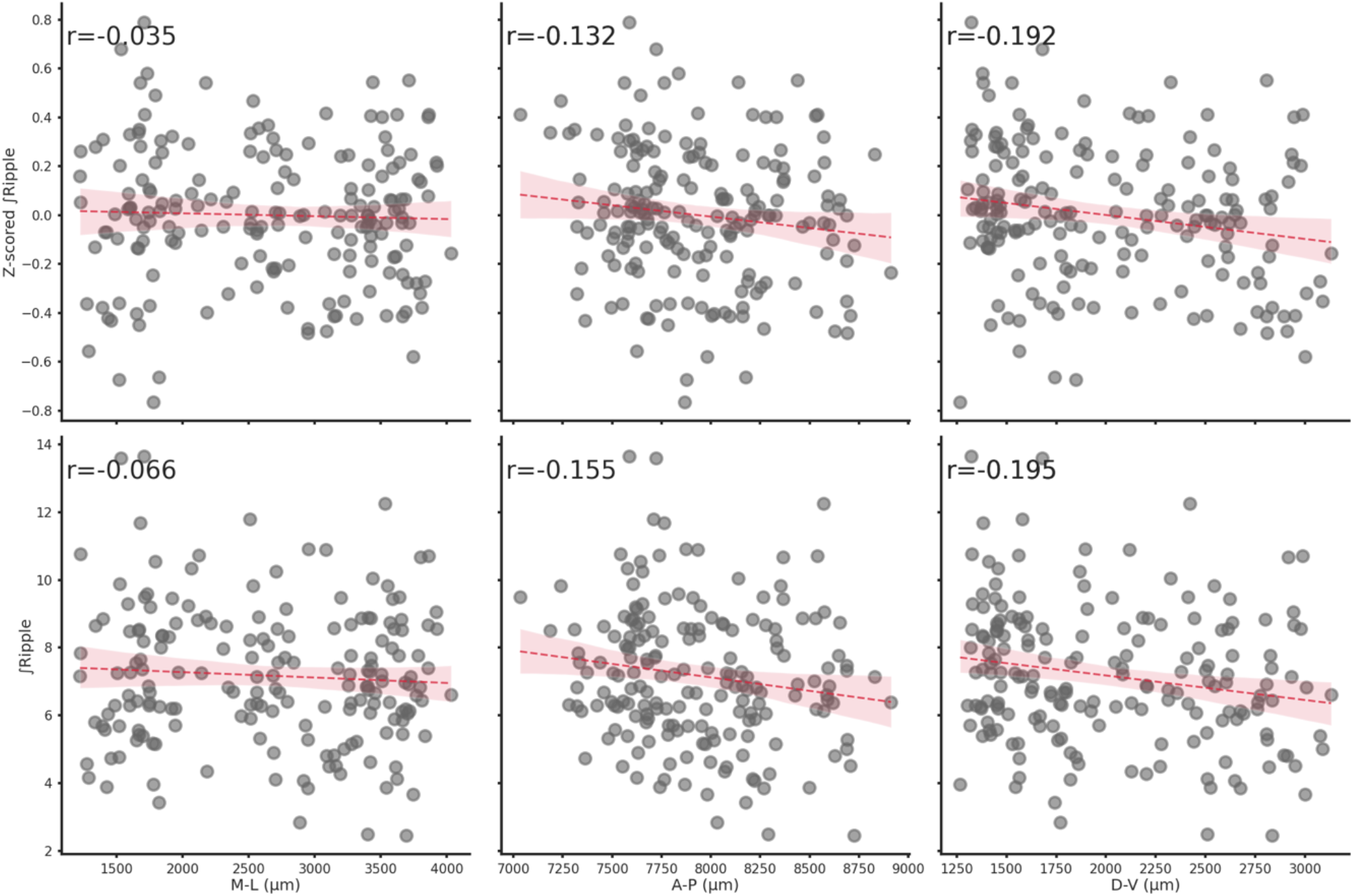
Spatial location does not influence ∫Ripple. Relationship between Z-scored ∫Ripple (top row) or ∫Ripple (bottom row) and each spatial axis (M-L, A-P or D-V). Spatial location has a negligible effect on ∫Ripple.

**Supplementary Figure 9.**
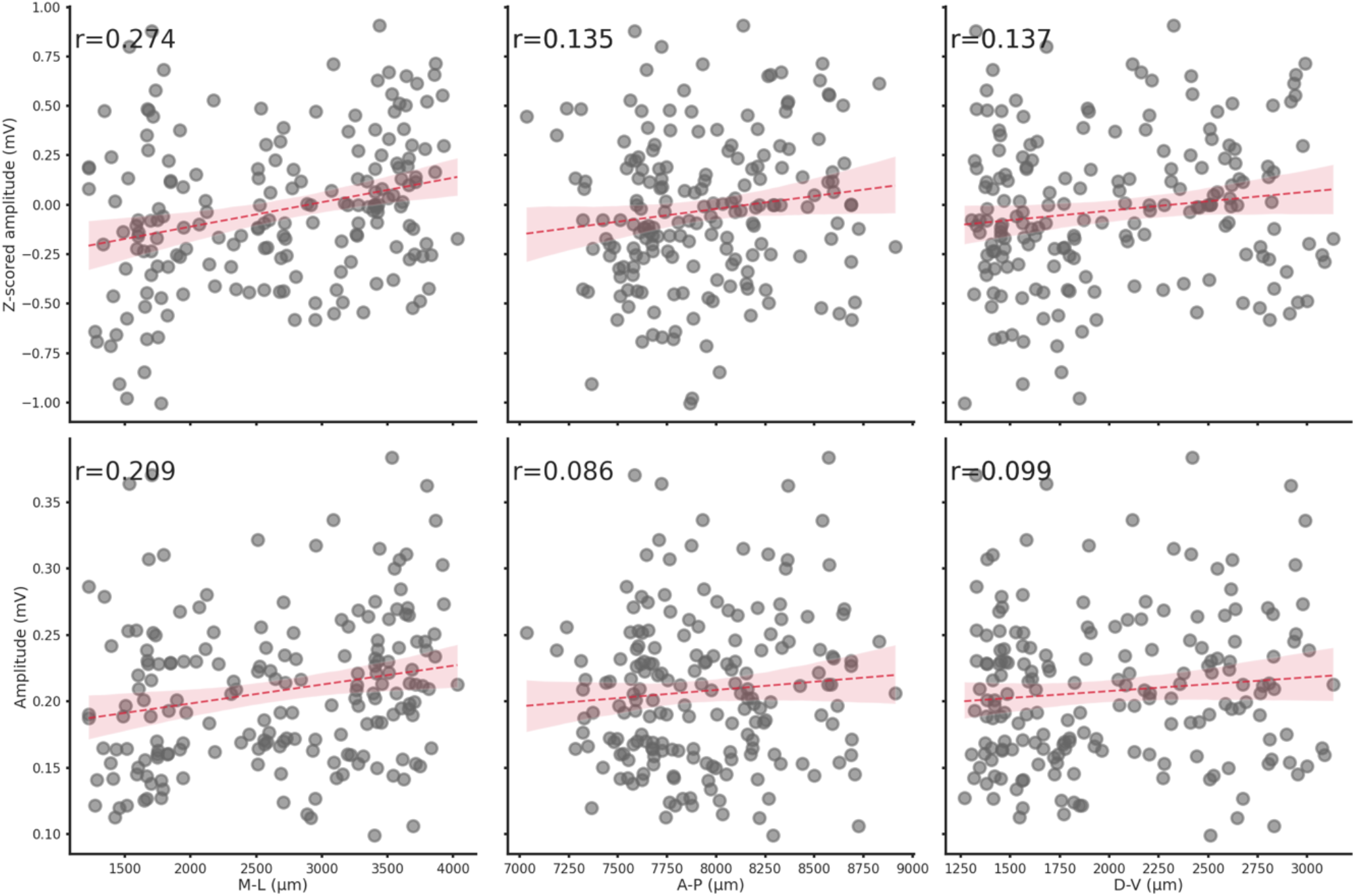
Spatial location does not influence ripple amplitude. Relationship between Z-scored amplitude (top row) or amplitude (bottom row) and each spatial axis (M-L, A-P or D-V). Spatial location has a negligible effect on ripple amplitude.

**Supplementary Figure 10.**
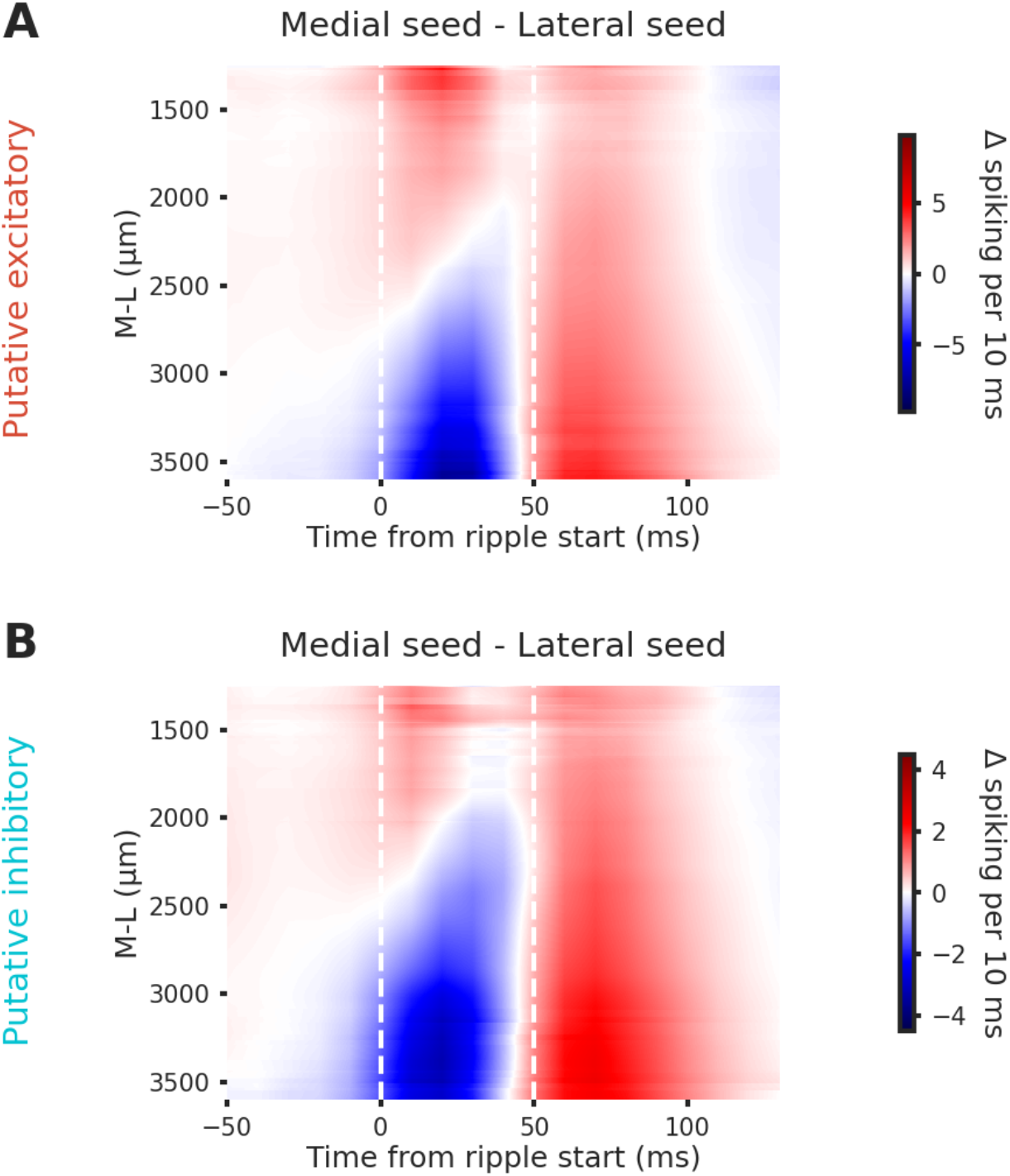
Putative excitatory and inhibitory neurons show similiar spiking patterns in lateral and medial ripples. Grand average of the differences between medial and lateral ripples induced spiking activity in putative excitatory (A) and inhibitory neurons (B).

**Supplementary Figure 11.**
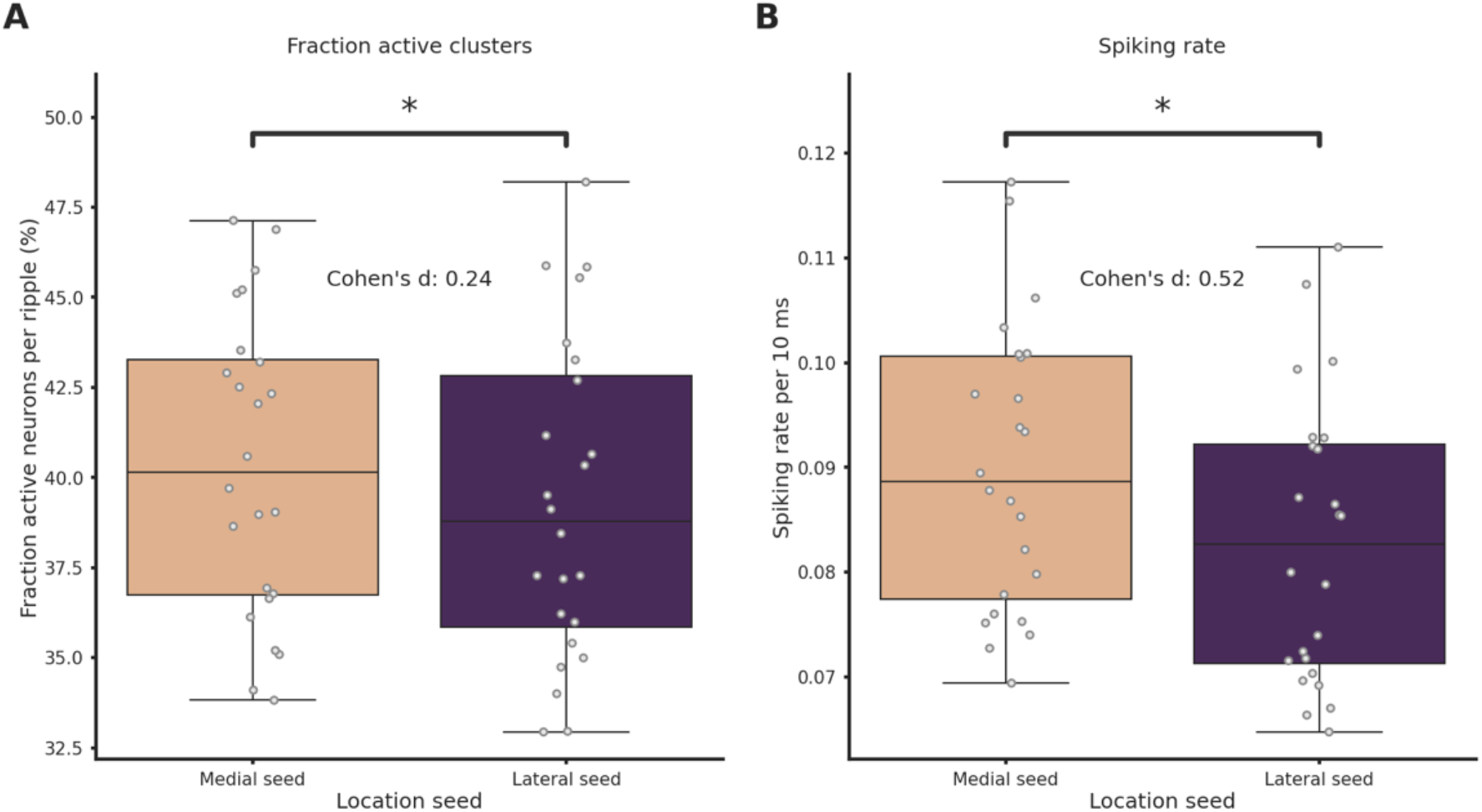
Spiking rate and fraction of active neurons are significantly higher in medial ripples. (A) Fraction of active neurons per ripple grouped by ripple seed location. (Medial seed=40.0±1.0%, lateral seed=39.0±1.0%, p-value=9.52e-05, Student’s t-test). (B) Average spiking rate grouped per ripple grouped by ripple seed location (Medial seed=9.0±0.0%, lateral seed=8.0±0.0%, p-value=5.20e-10, Student’s t-test). Asterisks mean p < 0.05, Student’s t-test.

**Supplementary Figure 12.**
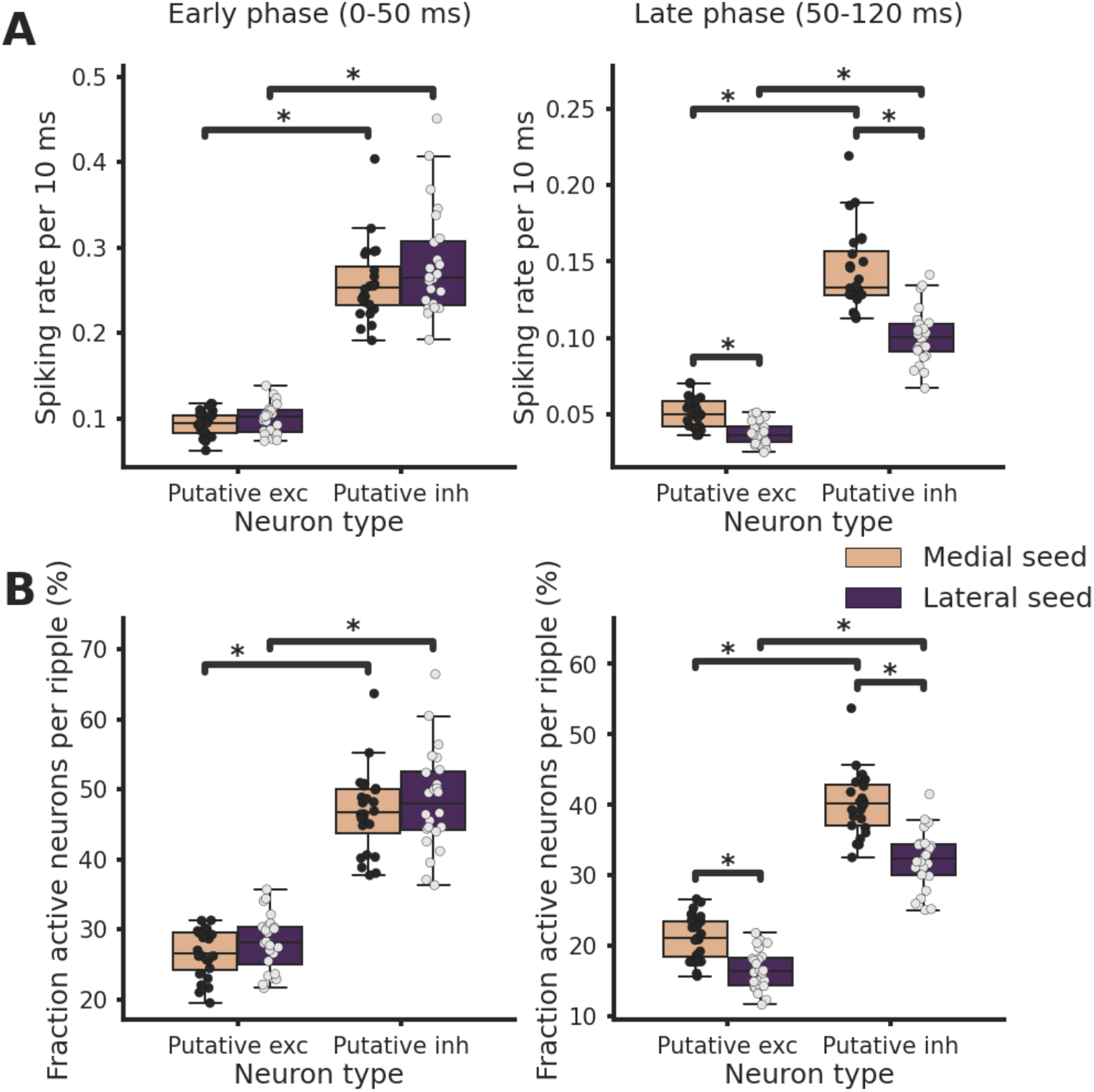
Spiking rate and fraction of active neurons are increased in the late phase post-ripple start in medial ripples both in putative excitatory and inhibitory neurons. (A) Average spiking rate in early (left) and late (right) phase post-ripple start grouped by ripple seed location and putative neuron identity. Asterisks mean p < 0.05, ANOVA with pairwise Tukey post-hoc test. (B) Fraction of active neurons per ripple in early (left) and late (right) phase post-ripple start grouped by ripple seed location and putative neuron identity. Asterisks mean p < 0.05, ANOVA with pairwise Tukey post-hoc test.

**Supplementary Figure 13.**
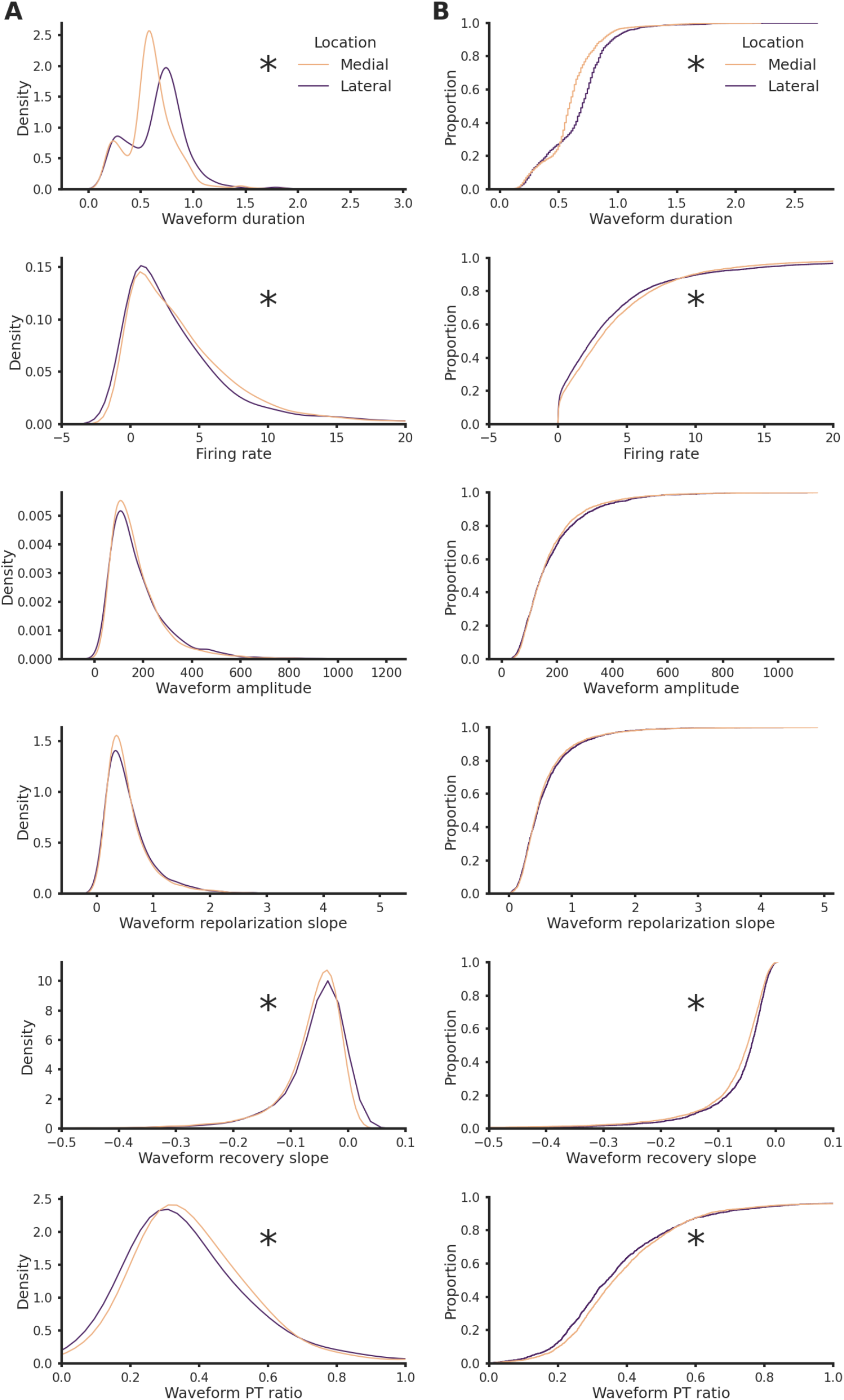
Units features in medial and lateral sections. (A) Kernel density estimate plot of waveform duration (p-value=1.64e-33), firing rate (p-value=6.41e-01), waveform amplitude (p-value=5.48e-01), waveform repolarization slope (p-value=4.09e-01), waveform recovery slope (p-value=1.13e-10) and waveform peak-through ratio (p-value=5.42e-05) grouped by hippocampal section.Asterisks mean p<0.05, Mann-Whitney U test. (B) Cumulative distribution plot of waveform duration (p-value=0.00e+00), firing rate (p-value=9.26e-03), waveform amplitude (p-value=9.09e-02), waveform repolarization slope (p-value=6.90e-02), waveform recovery slope (p-value=1.58e-10) and waveform peak-through ratio (p-value=2.27e-05) grouped by hippocampal section.Asterisks mean p < 0.05, Kolgomorov-Smirnov test.

## References

Bradfield, L. A., Leung, B. K., Boldt, S., Liang, S. & Balleine, B. W. 2020. Goal-directed actions transiently depend on dorsal hippocampus. Nature Neuroscience, 23, 1194–1197.

Buzsáki, G. 1989. Two-stage model of memory trace formation: a role for “noisy” brain states. Neuroscience, 31, 551–70.

Buzsáki, G. 2015. Hippocampal sharp wave-ripple: A cognitive biomarker for episodic memory and planning. Hippocampus, 25, 1073–188.

Carr, M. F., Jadhav, S. P. & Frank, L. M. 2011. Hippocampal replay in the awake state: a potential substrate for memory consolidation and retrieval. Nat Neurosci, 14, 147–53.

Claudi, F., Tyson, A. L., Petrucco, L., Margrie, T. W., Portugues, R. & Branco, T. 2021. Visualizing anatomically registered data with brainrender. eLife, 10, e65751.

Davidson, T. J., Kloosterman, F. & Wilson, M. A. 2009. Hippocampal replay of extended experience. Neuron, 63, 497–507.

Diba, K. & Buzsáki, G. 2007. Forward and reverse hippocampal place-cell sequences during ripples. Nat Neurosci, 10, 1241–2.

Diekelmann, S. & Born, J. 2010. The memory function of sleep. Nature Reviews Neuroscience, 11, 114–126.

Dragoi, G. & Tonegawa, S. 2011. Preplay of future place cell sequences by hippocampal cellular assemblies. Nature, 469, 397–401.

Fanselow, M. S. & Dong, H. W. 2010. Are the dorsal and ventral hippocampus functionally distinct structures? Neuron, 65, 7–19.

Fernández-Ruiz, A., Oliva, A., Fermino De Oliveira, E., Rocha-Almeida, F., Tingley, D. & Buzsáki, G. 2019. Long-duration hippocampal sharp wave ripples improve memory. Science (New York, N.Y.), 364, 1082–1086.

Foster, D. J. & Wilson, M. A. 2006. Reverse replay of behavioural sequences in hippocampal place cells during the awake state. Nature, 440, 680–3.

Gais, S., Albouy, G., Boly, M., Dang-Vu, T. T., Darsaud, A., Desseilles, M., Rauchs, G., Schabus, M., Sterpenich, V., Vandewalle, G., Maquet, P. & Peigneux, P. 2007. Sleep transforms the cerebral trace of declarative memories. Proc Natl Acad Sci U S A, 104, 18778–83.

Girardeau, G., Benchenane, K., Wiener, S. I., Buzsáki, G. & Zugaro, M. B. 2009. Selective suppression of hippocampal ripples impairs spatial memory. Nat Neurosci, 12, 1222–3.

Girardeau, G. & Zugaro, M. 2011. Hippocampal ripples and memory consolidation. Curr Opin Neurobiol, 21, 452–9.

Hock, B. J., Jr. & Bunsey, M. D. 1998. Differential effects of dorsal and ventral hippocampal lesions. The Journal of neuroscience: the official journal of the Society for Neuroscience, 18, 7027–7032.

Hulse, B. K., Moreaux, L. C., Lubenov, E. V. & Siapas, A. G. 2016. Membrane Potential Dynamics of CA1 Pyramidal Neurons during Hippocampal Ripples in Awake Mice. Neuron, 89, 800–13.

Jadhav, S. P., Kemere, C., German, P. W. & Frank, L. M. 2012. Awake hippocampal sharp-wave ripples support spatial memory. Science, 336, 1454–8.

Kheirbek, M. A., Drew, L. J., Burghardt, N. S., Costantini, D. O., Tannenholz, L., Ahmari, S. E., Zeng, H., Fenton, A. A. & Hen, R. 2013. Differential control of learning and anxiety along the dorsoventral axis of the dentate gyrus. Neuron, 77, 955–968.

Khodagholy, D., Gelinas, J. N. & Buzsáki, G. 2017. Learning-enhanced coupling between ripple oscillations in association cortices and hippocampus. Science, 358, 369–372.

Klinzing, J. G., Niethard, N. & Born, J. 2019. Mechanisms of systems memory consolidation during sleep. Nature Neuroscience, 22, 1598–1610.

Kumar, M. & Deshmukh, S. S. 2020. Differential propagation of ripples along the proximodistal and septotemporal axes of dorsal CA1 of rats. Hippocampus, 30, 970–986.

Maras, P. M., Molet, J., Chen, Y., Rice, C., Ji, S. G., Solodkin, A. & Baram, T. Z. 2014. Preferential loss of dorsal-hippocampus synapses underlies memory impairments provoked by short, multimodal stress. Mol Psychiatry, 19, 811–22.

Marr, D. 1971. Simple memory: a theory for archicortex. Philos Trans R Soc Lond B Biol Sci, 262, 23–81.

Mcclelland, J. L., Mcnaughton, B. L. & O’Reilly, R. C. 1995. Why there are complementary learning systems in the hippocampus and neocortex: insights from the successes and failures of connectionist models of learning and memory. Psychol Rev, 102, 419–457.

Mcglinchey, E. M. & Aston-Jones, G. 2018. Dorsal Hippocampus Drives Context-Induced Cocaine Seeking via Inputs to Lateral Septum. Neuropsychopharmacology, 43, 987–1000.

Moser, E., Moser, M. & Andersen, P. 1993. Spatial learning impairment parallels the magnitude of dorsal hippocampal lesions, but is hardly present following ventral lesions. The Journal of Neuroscience, 13, 3916–3925.

Moser, M. B. & Moser, E. I. 1998. Functional differentiation in the hippocampus. Hippocampus, 8, 608–19.

Moser, M. B., Moser, E. I., Forrest, E., Andersen, P. & Morris, R. G. 1995. Spatial learning with a minislab in the dorsal hippocampus. Proceedings of the National Academy of Sciences, 92, 9697–9701.

Ngo, H. V., Fell, J. & Staresina, B. 2020. Sleep spindles mediate hippocampal-neocortical coupling during long-duration ripples. Elife, 9.

Nitzan, N., Swanson, R., Schmitz, D. & Buzsáki, G. 2022. Brain-wide interactions during hippocampal sharp wave ripples. Proc Natl Acad Sci U S A, 119, e2200931119.

Noguchi, A., Huszár, R., Morikawa, S., Buzsáki, G. & Ikegaya, Y. 2022. Inhibition allocates spikes during hippocampal ripples. Nat Commun, 13, 1280.

Norman, Y., Yeagle, E. M., Khuvis, S., Harel, M., Mehta, A. D. & Malach, R. 2019. Hippocampal sharp-wave ripples linked to visual episodic recollection in humans. Science, 365, eaax1030.

Novitskaya, Y., Sara, S. J., Logothetis, N. K. & Eschenko, O. 2016. Ripple-triggered stimulation of the locus coeruleus during post-learning sleep disrupts ripple/spindle coupling and impairs memory consolidation. Learn Mem, 23, 238–48.

Patel, J., Schomburg, E. W., Berényi, A., Fujisawa, S. & Buzsáki, G. 2013. Local generation and propagation of ripples along the septotemporal axis of the hippocampus. J Neurosci, 33, 17029–41.

Pfeiffer, B. E. & Foster, D. J. 2015. PLACE CELLS. Autoassociative dynamics in the generation of sequences of hippocampal place cells. Science, 349, 180–3.

Qin, C., Bian, X.-L., Wu, H.-Y., Xian, J.-Y., Cai, C.-Y., Lin, Y.-H., Zhou, Y., Kou, X.-L., Chang, L., Luo, C.-X. & Zhu, D.-Y. 2020. Dorsal Hippocampus to Infralimbic Cortex Circuit is Essential for the Recall of Extinction Memory. Cerebral Cortex, 31, 1707–1718.

Ramirez-Villegas, J. F., Logothetis, N. K. & Besserve, M. 2015. Diversity of sharp-wave-ripple LFP signatures reveals differentiated brain-wide dynamical events. Proc Natl Acad Sci U S A, 112, E6379–87.

Rasch, B. & Born, J. 2007. Maintaining memories by reactivation. Curr Opin Neurobiol, 17, 698–703.

Roux, L., Hu, B., Eichler, R., Stark, E. & Buzsáki, G. 2017. Sharp wave ripples during learning stabilize the hippocampal spatial map. Nat Neurosci, 20, 845–853.

Siegle, J. H., Jia, X., Durand, S., Gale, S., Bennett, C., Graddis, N., Heller, G., Ramirez, T. K., Choi, H., Luviano, J. A., Groblewski, P. A., Ahmed, R., Arkhipov, A., Bernard, A., Billeh, Y. N., Brown, D., Buice, M. A., Cain, N., Caldejon, S., Casal, L., Cho, A., Chvilicek, M., Cox, T. C., Dai, K., Denman, D. J., De Vries, S. E. J., Dietzman, R., Esposito, L., Farrell, C., Feng, D., Galbraith, J., Garrett, M., Gelfand, E. C., Hancock, N., Harris, J. A., Howard, R., Hu, B., Hytnen, R., Iyer, R., Jessett, E., Johnson, K., Kato, I., Kiggins, J., Lambert, S., Lecoq, J., Ledochowitsch, P., Lee, J. H., Leon, A., Li, Y., Liang, E., Long, F., Mace, K., Melchior, J., Millman, D., Mollenkopf, T., Nayan, C., Ng, L., Ngo, K., Nguyen, T., Nicovich, P. R., North, K., Ocker, G. K., Ollerenshaw, D., Oliver, M., Pachitariu, M., Perkins, J., Reding, M., Reid, D., Robertson, M., Ronellenfitch, K., Seid, S., Slaughterbeck, C., Stoecklin, M., Sullivan, D., Sutton, B., Swapp, J., Thompson, C., Turner, K., Wakeman, W., Whitesell, J. D., Williams, D., Williford, A., Young, R., Zeng, H., Naylor, S., Phillips, J. W., Reid, R. C., Mihalas, S., Olsen, S. R. & Koch, C. 2021. Survey of spiking in the mouse visual system reveals functional hierarchy. Nature, 592, 86–92.

Sirota, A., Csicsvari, J., Buhl, D. & Buzsáki, G. 2003. Communication between neocortex and hippocampus during sleep in rodents. Proc Natl Acad Sci U S A, 100, 2065–9.

Sosa, M., Joo, H. R. & Frank, L. M. 2020. Dorsal and Ventral Hippocampal Sharp-Wave Ripples Activate Distinct Nucleus Accumbens Networks. Neuron, 105, 725-741.e8.

Steffenach, H. A., Witter, M., Moser, M. B. & Moser, E. I. 2005. Spatial memory in the rat requires the dorsolateral band of the entorhinal cortex. Neuron, 45, 301–13.

Strange, B. A., Witter, M. P., Lein, E. S. & Moser, E. I. 2014. Functional organization of the hippocampal longitudinal axis. Nat Rev Neurosci, 15, 655–69.

Takahashi, S. 2015. Episodic-like memory trace in awake replay of hippocampal place cell activity sequences. Elife, 4, e08105.

Tong, A. P. S., Vaz, A. P., Wittig, J. H., Inati, S. K. & Zaghloul, K. A. 2021. Ripples reflect a spectrum of synchronous spiking activity in human anterior temporal lobe. Elife, 10.

Tukker, J. J., Beed, P., Schmitz, D., Larkum, M. E. & Sachdev, R. N. S. 2020. Up and Down States and Memory Consolidation Across Somatosensory, Entorhinal, and Hippocampal Cortices. Front Syst Neurosci, 14, 22.

Ul Haq, R., Anderson, M. L., Hollnagel, J.-O., Worschech, F., Sherkheli, M. A., Behrens, C. J. & Heinemann, U. 2016. Serotonin dependent masking of hippocampal sharp wave ripples. Neuropharmacology, 101, 188–203.

Ul Haq, R., Liotta, A., Kovacs, R., Ræsler, A., Jarosch, M. J., Heinemann, U. & Behrens, C. J. 2012. Adrenergic modulation of sharp wave-ripple activity in rat hippocampal slices. Hippocampus, 22, 516–533.

Van Strien, N. M., Cappaert, N. L. & Witter, M. P. 2009. The anatomy of memory: an interactive overview of the parahippocampal-hippocampal network. Nat Rev Neurosci, 10, 272–82.

Vaz, A. P., Inati, S. K., Brunel, N. & Zaghloul, K. A. 2019. Coupled ripple oscillations between the medial temporal lobe and neocortex retrieve human memory. Science, 363, 975–978.

Vogel, J. W., La Joie, R., Grothe, M. J., Diaz-Papkovich, A., Doyle, A., Vachon-Presseau, E., Lepage, C., Vos De Wael, R., Thomas, R. A., Iturria-Medina, Y., Bernhardt, B., Rabinovici, G. D. & Evans, A. C. 2020. A molecular gradient along the longitudinal axis of the human hippocampus informs large-scale behavioral systems. Nat Commun, 11, 960.

Wang, D., Yau, H.-J., Broker, C., Tsou, J.-H., Bonci, A. & Ikemoto, S. 2015. Mesopontine median raphe regulates hippocampal ripple oscillation and memory consolidation. Nature neuroscience, 18.

Witter, M. P. 2007. Intrinsic and extrinsic wiring of CA3: indications for connectional heterogeneity. Learn Mem, 14, 705–13.

Xu, H., Baracskay, P., O’Neill, J. & Csicsvari, J. 2019. Assembly Responses of Hippocampal CA1 Place Cells Predict Learned Behavior in Goal-Directed Spatial Tasks on the Radial Eight-Arm Maze. Neuron, 101, 119-132.e4.

Ylinen, A., Bragin, A., Nádasdy, Z., Jandö, G., Szabö, I., Sik, A. & Buzsáki, G. 1995. Sharp wave-associated high-frequency oscillation (200 Hz) in the intact hippocampus: network and intracellular mechanisms. J Neurosci, 15, 30–46.

Zhang, Y., Cao, L., Varga, V., Jing, M., Karadas, M., Li, Y. & Buzsáki, G. 2021. Cholinergic suppression of hippocampal sharp-wave ripples impairs working memory. Proc Natl Acad Sci U S A, 118.

